# Origin of Eukaryotic Plasmalogen Biosynthesis by Horizontal Gene Transfer from Myxobacteria

**DOI:** 10.1101/2025.10.10.681575

**Authors:** Juan Manuel Trinidad-Barnech, Irene del Rey Navalón, Konstantina Mitsi, Antonio Joaquín Monera Girona, Sebastián R. Najle, S. Padmanabhan, Iñaki Ruiz-Trillo, Montserrat Elías-Arnanz

**Affiliations:** Laboratorio de Genómica Evolutiva, Sección Biomatemática, Facultad de Ciencias, Universidad de la República, Montevideo, Uruguay; Laboratorio de Bioinformática, Departamento de Genómica, Instituto de Investigaciones Biológicas Clemente Estable, MEC, Montevideo, Uruguay; Departamento de Genética y Microbiología, Área de Genética (Unidad Asociada al IQF-CSIC), Universidad de Murcia, 30100 Murcia, Spain; Institut de Biologia Evolutiva (CSIC-Universitat Pompeu Fabra), 08003 Barcelona, Spain; Instituto Español de Oceanografía, Centro Oceanográfico de Santander, Spain; Department of Biomedical Sciences, Faculty of Medicine and Health Sciences, Universitat Internacional de Catalunya (UIC), 08195 Sant Cugat del Vallès, Spain; Instituto de Química Física “Blas Cabrera”, CSIC (IQF-CSIC), 28006 Madrid, Spain; ICREA, Pg. Lluís Companys 23, 08010 Barcelona, Spain

**Keywords:** Plasmalogen, horizontal gene transfer, aerobic lipid biosynthesis, myxobacteria, PEDS1

## Abstract

Plasmalogens, a unique class of membrane lipids defined by a distinctive vinyl ether bond, are critical for human health, with their altered levels linked to various diseases. Despite their importance, their origin and evolutionary history remain enigmatic. Here, we uncover the evolutionary history of the aerobic plasmalogen biosynthesis pathway in eukaryotes, focusing on the four essential enzymes responsible for their formation. Through comprehensive phylogenetic analyses and experimental validation, we demonstrate a significant divide in plasmalogen synthesis capabilities across major eukaryotic lineages. Our study also suggests that the acquisition of these plasmalogen biosynthesis genes by an early eukaryotic ancestor was through horizontal gene transfer (HGT) from Myxobacteria. The findings yield insights into how HGT shapes metabolic pathways and illuminate a critical step in the genesis of eukaryotic cell complexity.

## Introduction

Lipids are crucial components of one of the defining features of cellular life, the membranes. Lipids determine membrane rigidity, fluidity and function, and serve as platforms for other crucial biomolecules like proteins and carbohydrates (1). One of the most abundant lipid classes are the glycerophospholipids or GPs (**Fig. 1A**). GPs have a glycerol backbone with a phosphate group, often bearing a head group such as ethanolamine or choline (1, 2). In bacteria and eukaryotes, this head group is at position *sn* (stereospecific numbering)-3, whereas in Archaea it is at position *sn*-1. Additionally, GPs also possess diverse acyl or alkyl sidechains at one or both remaining OH positions (1–3).

**Figure 1.**
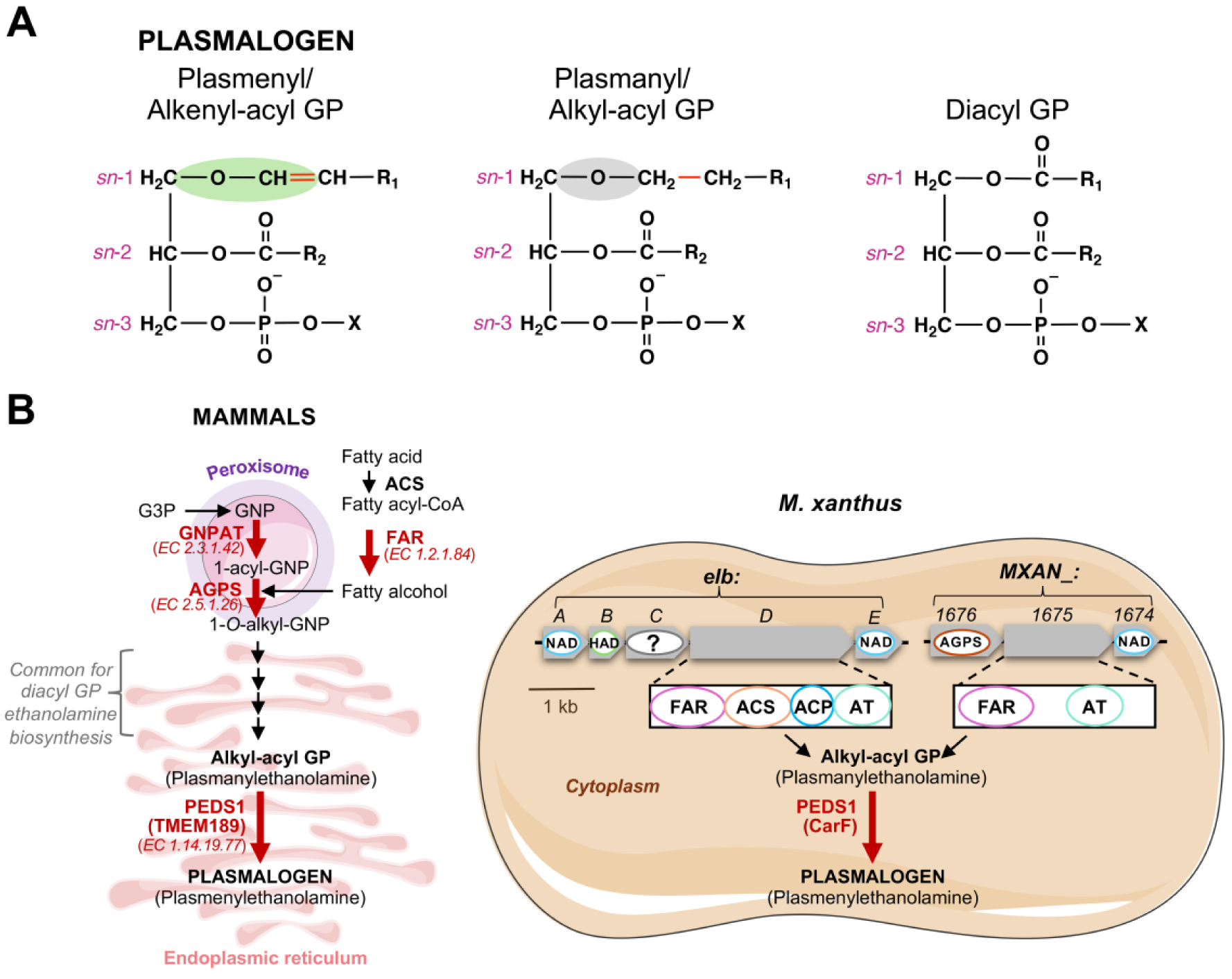
Plasmalogens and their aerobic biosynthesis pathway in mammals and in *M. xanthus*. (**A**) Chemical structures of plasmalogens (plasmenyl or alkenyl-acyl GPs), plasmanyl (or alkyl-acyl) GPs and diacyl GPs. The vinyl ether bond is shaded green, and the ether bond is shaded grey (the carbon-carbon bond that becomes desaturated by PEDS1 is marked in red). The *sn* positions are indicated, with the hydrocarbon chain at *sn*-1 represented as R_1_ and that at *sn*-2 as R_2_. X indicates the head group. **(B)** Left: Biosynthesis of plasmalogens in mammals. Signature enzymes of the pathway (FAR, GNPAT, AGPS and PEDS1) are indicated, with their Enzyme Comission numbers in brackets. For simplicity, steps in the biosynthesis pathway that are common for alkyl-acyl and diacyl GPs are not shown. G3P: glycerol 3-phosphate. Right: The two *M. xanthus* operons where *elbD* and *MXAN_1676* are located, with genes depicted as thick arrows and ovals inside them indicating that they encode for putative AGPS, FAR, AGPAT/GNPAT (AT), acyl-CoA synthase (ACS), acyl-carrier protein (ACP), NAD-dependent epimerase/dehydratase family protein (NAD), haloacid dehalogenase-like phosphohydrolase (HAD) functions. Gene deletions of *elbD* or *MXAN_1676* block alkyl and alkenyl ether lipid production via the corresponding biosynthetic pathway in *M. xanthus*.

A special type of GPs are the plasmalogens, which differ from diacyl GPs in that their *sn*-1 hydrocarbon chain is linked by a vinyl ether (1-*O-*alk-1′-enyl) bond, rather than the usual ester bond (**Fig. 1A**). This endows them with unique physical-chemical properties that can affect membrane fluidity, signaling and various biological functions (4–9). Plasmalogens can represent up to a fifth of the total human phospholipidome, and their deficiency correlates with various conditions, including rare diseases, cancer, neurodegenerative diseases, ferroptosis, responses to hypoxia, myeloid cell biology and inflammation (4–13). Nonetheless, the exact biological functions and modes of action of plasmalogens remain unclear.

Plasmalogens have a patchy distribution in the tree of life (6, 14). While they are present in diverse anaerobic and facultatively anaerobic bacteria (6, 15, 16), their presence in obligately aerobic bacteria is largely restricted to the phylum Myxococcota, where they have been most studied in the soil myxobacterium *Myxococcus xanthus* (7, 14, 17, 18). Among eukaryotes, plasmalogens have been reported in animals (Metazoa) and in a few protists but are conspicuously absent in plants and fungi (6, 14, 19–22). The multistep aerobic plasmalogen biosynthesis pathway (**Fig. 1B**), which differs considerably from the one that occurs in strict or facultative anaerobes (6, 15, 16), has been well-characterized in mammals (4–9) but is less studied in *M. xanthus* (7, 14, 17, 18). The mammalian aerobic pathway (**Fig. 1B**) begins in the peroxisome, where three key enzymes ensure formation of the early ether-bonded intermediate 1-*O*-alkyl-GNP (4–9). One is the peroxisomal membrane-anchored enzyme fatty acyl-CoA reductase 1 or FAR1 (or its isozyme FAR2), which catalyzes reduction of saturated or unsaturated fatty acyl-CoA to fatty alcohol. The second is glycerone phosphate *O*-acyltransferase or GNPAT, which converts glycolysis-derived glycerone phosphate (GNP) to 1-acyl-GNP. GNPAT acts as a complex with the third enzyme, alkylglyceronephosphate synthase (AGPS), which generates 1-*O*-alkyl-GNP that is shuttled to the endoplasmic reticulum. Various steps in the endoplasmic reticulum, common to those utilized for the biosynthesis of diacyl GPs with ethanolamine as the head group, generate the plasmanyl or alkyl-acyl GPs (plasmanylethanolamine), which plasmanylethanolamine desaturase or PEDS1 converts into plasmalogens (plasmenyl or alkenyl-acyl GPs; **Fig. 1B**). While FAR1, GNPAT, and AGPS are defining enzymes for plasmanyl GP biosynthesis, PEDS1 is the crucial desaturase that transforms plasmanyl GPs into plasmalogens.

The actual identity of PEDS1 was only recently unmasked and shown to correspond to *M. xanthus* protein CarF and its various metazoan homologs (14, 23, 24). Identifying PEDS1 resolved a key gap in the complex well-charted multistep pathway for aerobic biosynthesis of plasmalogens in mammals and suggested parallels with plasmalogen biosynthesis in *M. xanthus*, where two pathways coexist for plasmanyl GP production (**Fig. 1B**) (7, 14, 18). A key player in one pathway is ElbD, a large multidomain protein with annotated FAR and acylglycerolphosphate acyltransferase (AGPAT/GNPAT) modules, encoded in an operon with four other genes of still undefined roles, and with no apparent AGPS candidate (7, 14, 18). The other pathway involves a three-gene operon encoding: a putative AGPS (old locus tag MXAN_1676; current locus tag MXAN_RS08130) with ∼42% sequence identity to its human counterpart and experimentally demonstrated to be implicated in ether lipid biosynthesis in *M. xanthus* (14); a two-domain protein with predicted FAR and GNPAT modules (old locus tag MXAN_1675; current locus tag MXAN_RS08125); and a possible NAD-dependent epimerase/dehydratase family protein (old locus tag MXAN_1674; current locus tag MXAN_RS08120) (7, 14, 18, 25). Thus, besides CarF/PEDS1, which exhibits a remarkable 46% identity with human PEDS1, *M. xanthus* has potential FAR, GNPAT and AGPS activities, as in the aerobic mammalian plasmalogen biosynthesis pathway.

Despite the importance of plasmalogens in humans, the origin and evolution of their biosynthesis in eukaryotes remain unknown. Understanding this is highly relevant given the compelling issues of membrane evolution, the Bacteria/Eukarya versus Archaea lipid divide, and eukaryogenesis. In this regard, the intriguing distribution of plasmalogens across the tree of life, together with the discovery of the remarkable conservation of PEDS1 and possibly other key players across a vast evolutionary distance to aerobic myxobacteria, is particularly propitious. Yet, only a few studies have tried to shed some light into this question. One study demonstrated that *M. xanthus* CarF and its human, mouse, zebrafish, fruit fly and nematode homologs (denoted TMEM189), as well as those from *Leptospira* (bacterial phylum Spirochaetota), all correspond to PEDS1; but those in plants or Alphaproteobacteria were not functionally equivalent to PEDS1 (14). A more recent study (22) identified PEDS1 homologs in some members of the Holozoa clade (26, 27), the group that includes animals and their closest unicellular relatives, some genera of Amoebozoa and Excavata, and even in a few fungi within Ascomycetes. However, the taxon sampling used in these studies was limited, largely constraining the potential interpretations.

In this study, we aimed to investigate the origin and evolution of the aerobic plasmalogen biosynthesis pathway in eukaryotes. To this end, we first revised the distribution of the key enzymes for plasmalogen biosynthesis by examining the presence of PEDS1 homologs and its co-existence with FAR, GNPAT and AGPS in eukaryotes, using a broad taxon and up-to-date eukaryotic proteomic dataset. PEDS1 homologs were found in Discoba and Amorphea clades (where they often co-occur with FAR, GNPAT and AGPS) but not in Metamonada (which also lack homologs to the other three enzymes) or Diaphoretickes (which often harbor the divergent and typically plant-associated Fatty Acid Desaturase 4 or FAD4). We experimentally validated this split in plasmalogen-biosynthesis capacity across eukaryotic supergroups by testing the plasmanylethanolamine desaturase activity of PEDS1 homologs in phylogenetically key organisms that represent pivotal branches of the eukaryotic tree of life and by confirming the presence of plasmalogens in a subset of these organisms. We implemented a phylogenetic approach to reconstruct the origin and evolutionary history of plasmalogen biosynthetic pathway and explored the subsequent evolutionary dynamics that have shaped the patchy distribution of plasmalogen biosynthesis across extant eukaryotic lineages. The findings presented herein illuminate a critical step in the evolution of eukaryotic cell complexity and provide insights into the role of HGT in shaping metabolic pathways.

## Results

### Distribution of plasmalogen biosynthesis pathway in eukaryotes

To unravel the evolution of the plasmalogen biosynthesis pathway in eukaryotes, we first performed an extensive search in a dataset of 376 high-quality genomes encompassing the whole breadth of the known eukaryotic diversity (see **Materials and Methods**) to map the distribution of FAR, GNPAT, AGPS and PEDS1. For improved detection of distant homologs, we followed an iterative search algorithm that accounts for genome variations over evolutionary time and assigns higher rates to conserved positions when evaluating the hits. This approach allows the identification of functionally conserved regions of proteins, even when overall sequence similarity is low (see **Materials and Methods**). Mapping the distribution of the four key enzymes involved in the biosynthesis of plasmalogens would not only provide valuable insights into the evolutionary origin of the metabolic pathway in eukaryotes but also into which groups are likely capable of synthesizing this special type of GPs. Detection of FAR, GNPAT and AGPS would suggest an ability to produce plasmalogen precursors, and the additional presence of PEDS1 would warrant plasmalogen synthesis.

#### Discoba

Within the Discoba clade, the plasmalogen biosynthesis pathway exhibited significant variation among the five groups analyzed (**Fig. 2A** and **Supplementary Tables S1-S5**). Thus, Heterolobosea and Kinetoplastea generally have PEDS1, whereas Jakobida, Euglenida, and Diplonemea lack this enzyme. Our analysis detected GNPAT hits in Heterolobosea, Kinetoplastea and Euglenida that, except for the one in *Angomonas deanei* (Kinetoplastea), had a large N-terminal region (∼650 amino acids) unassigned to any known domain and absent in mammalian GNPAT homologs (whose total size is typically ∼680 amino acids). Such an extension had been reported previously in the Trypanosomatidae family (28, 29) but no precise function could be assigned to it. Given that *M. xanthus* GNPAT in MXAN_1675 has an N-terminal domain with predicted FAR activity and that the *Tetrahymena thermophila* (Ciliophora) GNPAT was experimentally shown to occur as a C-terminal fusion to FAR in a single bifunctional 1140-residue FARAT protein (30), we explored the possibility that the large GNPAT hits were really FARAT. To address this, we used protein structure comparison, which has proven to be a sensitive alternative for protein domain identification and homology detection in Discoba (31, 32). Effectively, by using this approach, we could assign all the detected GNPAT with these long N-terminal extensions as FARAT, thus unmasking its presence in Heterolobosea, Euglenida and Kinetoplastea (**Supplementary Fig. S1** and **Supplementary Table S3**) (see **Materials and Methods**). In addition, we detected a second copy of FAR, as an independent protein, in *Euglena gracilis* and in the Trypanosomatidae *Strigomonas culicis* and *Vickermania ingenoplastis* (**Supplementary Table S4**). Finally, we observed AGPS in all groups except Jacobida, and always co-occurring with the other enzymes except in *Diplonema papillatum* (the only Diplonemea analyzed), where we could identify only AGPS (**Supplementary Table S5**).

**Figure 2.**
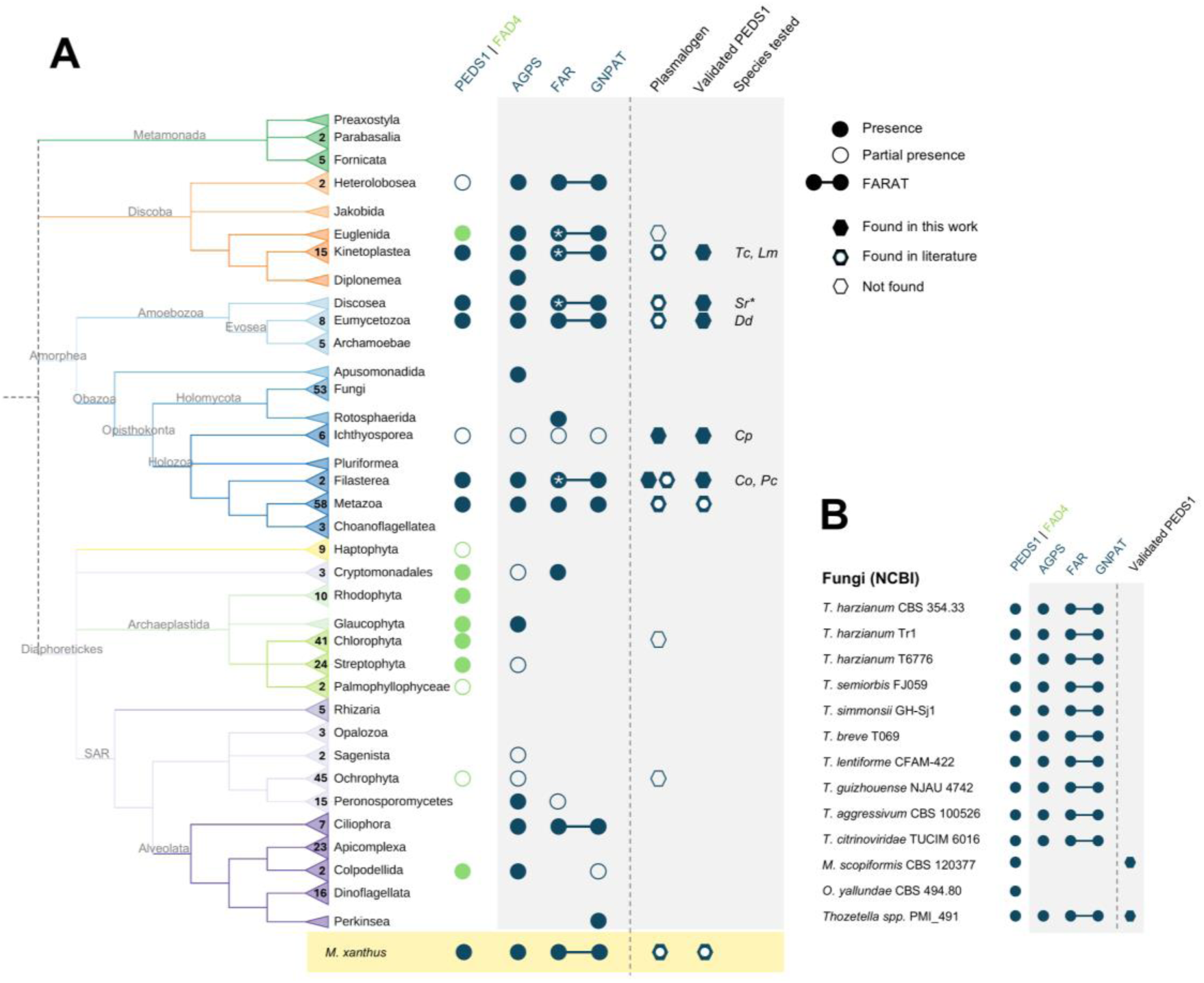
Distribution of plasmalogen pathway genes across major Eukaryotic clades. (**A**) A schematic phylogenetic tree illustrating the presence of key enzymes in the plasmalogen biosynthesis pathway (PEDS1, AGPS, FAR and GNPAT) across diverse eukaryotic lineages. The tree is color-coded by supergroup, with numbers in each triangular branch indicating how many genomes were analyzed and triangles without numbers representing clades with a single genome. Open or filled circles indicate, respectively, that more than one-third or two-thirds of species in the clade have the indicated enzyme. FAR and GNPAT connected by a line indicates a bifunctional FAR-GNPAT fused FARAT protein, and the FAR protein/domain with an asterisk indicates FAR presence also as an independent protein. In the “Plasmalogen” column: filled hexagons or with a circle indicate, respectively, that plasmalogens were detected in this study or in other studies; outline-only hexagons indicate that plasmalogens were not detected in lipid analyses. In the “Validated PEDS1” column: hexagons mark cases where PEDS1 function was confirmed by heterologous complementation in *M. xanthus* (filled hexagon, validated in this study; with a circle, in other studies). The final column lists species whose PEDS1 was tested in this study: *Trypanosoma cruzi* (*Tc*), *Leishmania major* (*Lm*), *Stygamoeba regulata* (*Sr*), *Dictyostelium discoideum* (*Dd*), *Chromosphaera perkinsii* (*Cp*), *Capsaspora owczarzaki* (*Co*), and *Pigoraptor chileana* (*Pc*). The asterisk next to *Sr* indicates that PEDS1 was identified through transcriptomic data. The tree was obtained from EukProt (33). Phylogenetic positions inferred from (27, 34, 35). (**B**) The small subset of fungal species where PEDS1 was found in BLASTP analyses and the presence in these species of the other enzymes in the plasmalogen biosynthesis pathway. Symbols as in (A).

While the genomes of both species analyzed within Heterolobosea, *Naegleria gruberi* and *Neovahlkampfia damariscottae*, encode AGPS and FARAT, only *N. gruberi* additionally harbors PEDS1, suggesting that plasmalogen biosynthesis is restricted to the latter. In Kinetoplastea, a group that is highly divergent from Heterolobosea, the pattern of occurrence was found to be complex, with ten out of fifteen taxa containing PEDS1, seven having all the enzymes and two (*Trypanosoma vivax* and *Perkinsela spp*.) lacking them entirely. The finding that homologs to the four enzymes occur in several species is consistent with previous experimental findings showing the presence of plasmalogens in various kinetoplastid taxa, including *Trypanosoma* (21, 36, 37), *Strigomonas* (38), *Crithidia* (37), and *Leishmania* (37, 39). Notably, species lacking PEDS1, such as *Bodo saltans* and *Phytomonas sp. Hart1* have the other three enzymes, suggesting a secondary loss of PEDS1 that would entail the ability to synthesize only the plasmanyl GPs.

The Euglenida class, represented by *E. gracilis*, is unique among Discoba in having FAD4 instead of PEDS1. FAD4 emerges in PEDS1 queries due to their sequence similarity but it is not functionally equivalent to PEDS1. This divergent lipid desaturase was first described in *Arabidopsis thaliana* as a chloroplast membrane-associated enzyme (40–42), in contrast to PEDS1 that localizes to the endoplasmic reticulum (14). Unlike PEDS1, FAD4 generates a *trans* double bond between carbon 3 and 4 (counting from the carboxyl end) of palmitic acid esterified at position *sn*-2 of phosphatidylglycerol (where the phosphate head group is also a glycerol) to generate a GP typically found in the thylakoid membranes throughout the plant kingdom, from green algae to different land plants (41). Lack of PEDS1 in *E. gracilis* is consistent with previous lipid analyses detecting plasmanyl GPs but no plasmalogens in this organism (37, 38, 43).

#### Amorphea

The Amorphea supergroup was also found to exhibit a heterogeneous distribution of the plasmalogen biosynthesis pathway (**Fig. 2A** and **Supplementary Tables S1-S5**), indicative of a complex evolutionary history shaped by differential gene retention. Within the Amoebozoa clade we examined two phyla: Discosea and Evosea (comprising Eumycetozoa and Archamoebae). In the sole representative analyzed of Discosea, *Acanthamoeba castellanii*, we found not only AGPS and FARAT (as well as FAR) but also PEDS1, consistent with the finding of plasmalogens in this species (37). Within the Evosea, the complete pathway was ubiquitously detected in the genomes of the Eumycetozoa (slime molds), with the single exception of *Synstelium polycarpum*, which lacks PEDS1. By contrast, no evidence of the plasmalogen pathway was found in Archamoebae, the sister lineage to Eumycetozoa, suggesting that the entire pathway was secondarily lost in this group. Similarly to Discoba, FAR and GNPAT were recurrently found as FARAT in Amoebozoa, but in the later we could reliably identify the FAR domain using PSI-BLAST and InterProScan, which made its detection straightforward. A FARAT protein among Amebozoa had only been previously described in *Dictyostelium discoideum* that, consistent with having a complete functional pathway, was shown to produce plasmalogens (44, 45). Only one enzyme of the pathway, AGPS, was detected in the Apusomonads *Thecamonas trahens*.

Among Holomycota, we found the entire pathway absent in Fungi but we report the presence of AGPS and FAR homologs in their closest relative, *Fonticula alba* (Rotosphaerida). Fungi are of particular interest because our analysis of 53 representative fungal genomes indicated a complete loss of the pathway (with two exceptions of AGPS present; **Supplementary Table S5**), consistent with the established notion that Fungi do not produce plasmalogens (6, 46). However, a recent report based on BLASTP analyses against the NCBI database suggested the presence of PEDS1 in three fungal species: *Thozetella* sp. (KAH8900397), *Trichoderma semiorbis* (KAH0527355), and *Mollisia scopiformis* (XP_018071845) (22). To further investigate the presence of PEDS1 in fungal taxa, we analyzed the gene structure and synteny of fungal PEDS1 homologs that we retrieved with BLASTP against NCBI nr database. Apart from the previously referred, we also found PEDS1 homologs in the genomes of *Oculimacula yallundae*, and eight species of the *Trichoderma* genus (**Fig. 2B**, **Supplementary Fig. S2** and **Supplementary Table S6**), all listed as monoisolate in NCBI. A search for AGPS, FAR and GNPAT in these fungal genomes detected none of these enzymes in *O. yallundae* and *M. scopiformis* but revealed their presence in *Thozetella* sp. and in all *Trichoderma* species carrying PEDS1, where the enzymes are generally encoded by neighboring genes (**Fig. 2B, Supplementary Fig. S2** and **Supplementary Table S6**). Interestingly, FAR and GNPAT were also found fused as FARAT in these fungal species, with *Thozetella* sp. carrying two copies of FARAT. The restricted occurrence of plasmalogen biosynthesis enzymes in a limited number of Fungi may be explained by two potential evolutionary scenarios, which will be discussed in a later section.

Among the Holozoa, the group that includes Metazoa and their closest unicellular relatives, we report the presence of the full plasmalogen pathway in the filasterean species *Pigoraptor chileana* and we confirm it in *Capsaspora owczarzaki* (22). Both species not only had FARAT but also an independent FAR copy. Out of six Ichthyosporean genomes analyzed, the full pathway was only detected in *Chromosphaera perkinsii*, which, instead of FARAT, carries separate GNPAT and FAR proteins. Except for AGPS in *Creolimax fragrantissima*, no evidence of the pathway was found in the remaining ichthyosporean species. Similarly, the pathway was absent from Corallochytrea (also known as Pluriformea) and Choanoflagellatea, the closest unicellular relatives of animals. In Metazoa, where the presence of plasmalogens has been widely reported (13, 14, 37, 47–50), the complete plasmalogen pathway was found in ∼70% of the 58 genomes analyzed, including all early-branching metazoan lineages. Interestingly, PEDS1 was absent from Metazoa with a parasitic lifestyle (∼10% of analyzed metazoan genomes), including *Acyrthosiphon pisum, Ixodes scapularis, Lepeophtheirus salmonids, Petromyzon marinus* and the two flatworm representatives in our dataset, *Schistosoma mansoni* and *Taenia solium*. Whereas *A. pisum* and *I. scapularis* contain all the genes except PEDS1, the two flatworms lack them entirely, and the other parasites contain only a partial set. Notably, we did not identify any instance of FARAT among Metazoa, where GNPAT sequences typically ranged between ∼600 and ∼750 amino acids (**Supplementary Table S2**). The GNPAT from the jellyfish *Sanderia malayensis* (1248 amino acids) had an N-terminal extension but with no detectable FAR domain based on BLASTP and InterProScan analyses, or structural predictions. Thus, independent genes for GNPAT and FAR are the norm in Metazoa, contrasting with the two being predominantly present as FARAT in other Amorphea groups.

#### Diaphoretickes

PEDS1 was not detected among Diaphoretickes (**Fig. 2A and Supplementary Table S1**) and the other enzymes were rarely found together in a genome or group (**Fig. 2A and Supplementary Tables S2 to S5**). One notable exception is the phylum Ciliophora (within the SAR group), where we could identify AGPS co-occurring with FAR and GNPAT, mostly fused as FARAT, in six out of the seven ciliophoran genomes analyzed (**Supplementary Tables S2 and S3**, and **Supplementary Fig. S3**). Thus, all genes identified initially as GNPAT, except for one in *Paramecium tetraurelia* and another in *Pseudocohnilembus persalinus*, were actually FARAT. PSI-BLAST was sufficient to identify at least one FARAT (as a GNPAT or FAR hit) in *P. tetraurelia* (with two FARATs), in *Stylonychia lemnae* (with three FARATs) and in *T. thermophila* (with a single FARAT). Overall, these results imply the ability of Ciliophora to produce plasmanyl GPs but not plasmalogens, in agreement with lipid analyses reported for *T. thermophila* (30).

The absence of PEDS1 in Diaphoretickes contrasts with the presence of FAD4 in several groups. As mentioned before, FAD4 is also a desaturase retrieved in the search for PEDS1 homologs, but its function is unrelated to plasmalogen biosynthesis (14, 41). Among the nine Haptophyta species analyzed, FAD4 was detected in three (*Diacronema lutheri, Emiliania huxleyi* and *Phaeocystis antarctica*), all of which lacked not only PEDS1 but also AGPS, GNPAT, and FAR. FAD4 was also detected in Cryptomonadales, including *Guillardia theta* and *Hanusia phi*, where AGPS, GNPAT and FAR showed a sporadic distribution (**Fig. 2A** and **Supplementary Tables S1, S2, S4 and S5**). In contrast, FAD4 was prevalent throughout Archaeplastida, encompassing red algae, green algae and plants (**Fig. 2A** and **Supplementary Table S1**). In Rhodophyta, eight of the ten species analyzed contained FAD4, three of which also had AGPS. In Glaucophyta, represented by *Cyanophora paradoxa*, FAD4 co-occurred with AGPS. Within Chloroplastida, comprising 67 genomes from Chlorophyta, Streptophyta and Palmophyllophyceae, FAD4 was detected in ∼80% of the species. In most of these, AGPS and/or FAR were absent, except for six genomes where both were found together with FAD4 (**Fig. 2A** and **Supplementary Tables S1, S2, S4 and S5**). Finally, within the SAR clade, FAD4 was restricted to Ochrophyta, where it was present in ∼64% of 45 genomes analyzed, and Colpodellida, where both representative species had FAD4 (**Fig. 2A** and **Supplementary Table S1**). By contrast, AGPS, GNPAT and FAR displayed a patchy distribution across SAR, except in Ciliophora, where, as described above, neither FAD4 nor PEDS1 were present (**Fig. 2A** and **Supplementary Tables S1, S2, S4 and S5**).

### Experimental analysis of selected PEDS1 homologs

Given the patchy distribution of PEDS1 in the tree of eukaryotes (**Fig. 2**), we wondered whether hits retrieved bioinformatically conserve the plasmanylethanolamine desaturase function. Therefore, we aimed to verify their enzymatic activity and the capacity for plasmalogen production in these organisms by testing selected PEDS1 homologs from representative taxa. PEDS1 homologs from various Metazoa have already been experimentally tested in previous studies (13, 14, 37, 47–50), so we focused on other taxonomic groups to provide a broad overview of PEDS1 functionality across key eukaryotic lineages. In particular, we chose the following (**Supplementary Fig. S4A** and **Supplementary Table S7**): *Capsaspora owczarzaki* and *Pigoraptor chileana*, in the Filasterea group, whose members are among the most closely related unicellular relatives of Metazoa (51); *Chromosphaera perkinsii* (Ichthyosporea); *D. discoideum* (Eumycetozoa) and *Stygamoeba regulata* (given the limited representation in Discosea), both in the major Amoebozoa taxonomic group; *Trypanosoma cruzi* and *Leishmania major*, both in Kinetoplastea.

To test for PEDS1 function we performed heterologous complementation experiments in *M. xanthus*, where plasmalogens are specifically required to induce a photoprotective carotenogenic response that causes cells to turn from yellow in the dark to orange in the light (7, 14, 52). The *M. xanthus* plasmalogen MxVEPE (1-*O*-(13-methyl-1-*Z*-tetradecenyl)-2-*O*-(13-methyltetradecanoyl)-glycero-3-phosphatidylethanolamine) differs from the plasmalogens produced in eukaryotes in having branched hydrocarbon chains (iso15:0 or i15:0) at *sn*-1 and *sn*-2, rather than the usual straight chains. Nonetheless, functional eukaryotic PEDS1 homologs have been shown to generate MxVEPE when expressed in the *carF* deleted (Δ*carF*) *M. xanthus* strain, even if they produce different plasmalogens in their natural organism (14). Thus, complementation can be monitored by the light-induced colony color change and by lipid analysis, with MxVEPE detected as the iso15:0 dimethylacetal (i15:0 DMA) derivative by fatty acid methyl ester gas chromatography-mass spectrometry (FAME GC-MS) analysis (14). The *M. xanthus* strains expressing each selected PEDS1 homolog (N-terminally FLAG-tagged for immunoblot analysis) in the Δ*carF* genetic background produced MxVEPE, albeit in different amounts (**Fig. 3A**). This is likely due to the observed differences in protein levels (**Fig. 3A**), together with possible intrinsic variations in activity and substrate preferences. The orange color under light correlated with MxVEPE levels produced by each homolog, and the color intensity in strains expressing homologs from *C. perkinsii*, *P. chileana* and *T. cruzi* closely matched that for *M. xanthus* CarF (**Fig. 3A**). Our data therefore confirm PEDS1 activity for the various homologs tested, which cover diverse taxonomic groups.

**Figure 3.**
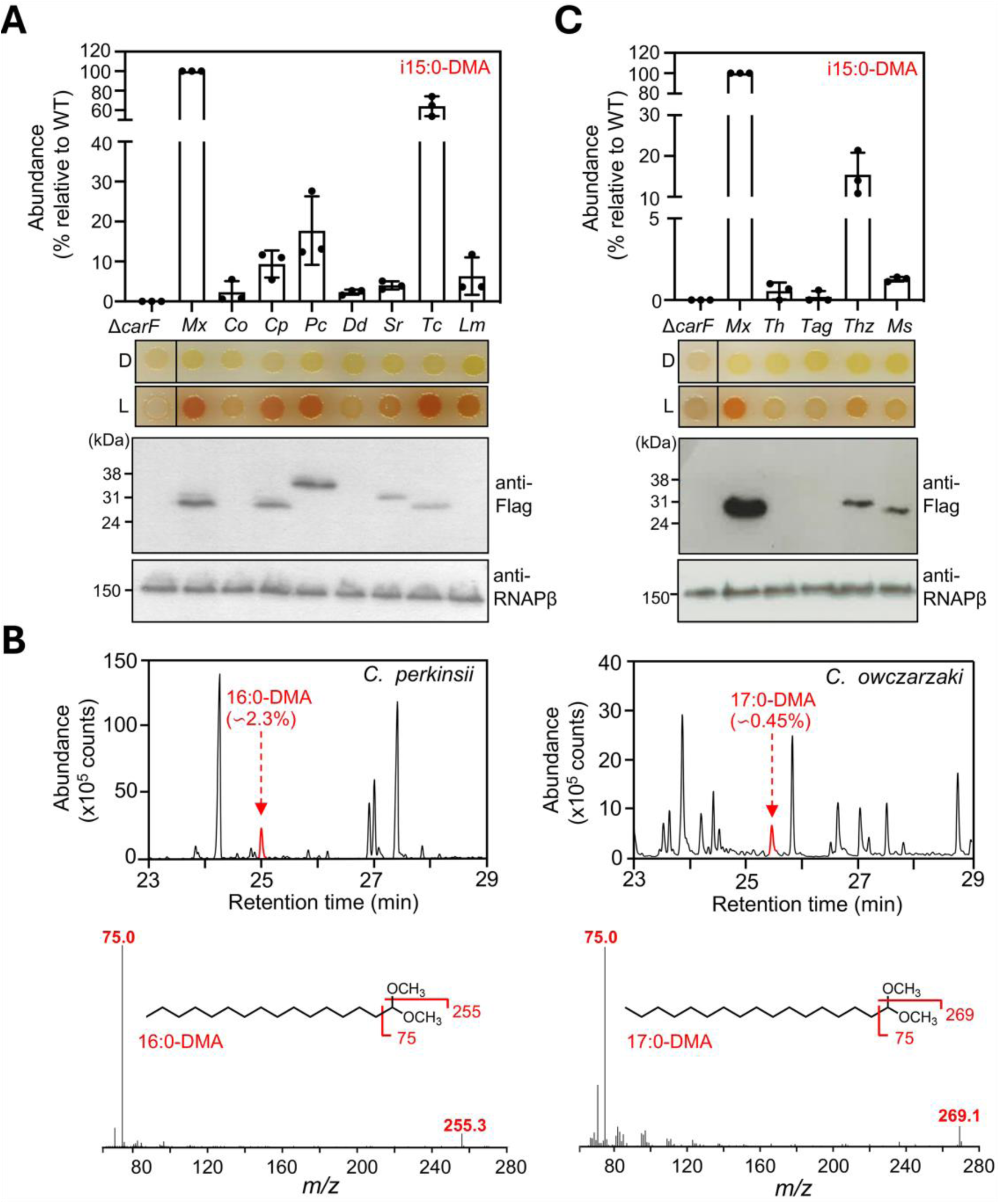
Functional validation of selected PEDS1 homologs. (**A**) Complementation analyses using the *M. xanthus* Δ*carF* strain expressing *M. xanthus* CarF (*Mx*) or homologs from *C. owczarzaki* (*Co*), *C. perkinsii* (*Cp*), *P. chileana* (*Pc*), *D. discoideum* (*Dd*), *S. regulata* (*Sr*), *T. cruzi* (*Tc*), and *L. major* (*Lm*) (all FLAG-tagged for immunoblot analysis). Top: Abundance of i15:0-DMA relative to 100% for the strain expressing *M. xanthus* CarF. Values are mean ± SD (n = 3 biological replicates). Middle: Light-induced colony-color assay for the corresponding strains above. Bottom: Immunoblot of the corresponding cell extracts probed with monoclonal anti-FLAG antibodies and, as loading control, polyclonal antibodies against the β subunit of RNAP. (**B**) Representative FAME GC-MS chromatogram (relevant section shown) of total lipid extracts (top) from *C. perkinsii* and *C. owczarzaki*. Electron impact mass spectrum and diagnostic *m/z* peaks (below) in FAME GC-MS analysis for the characteristic 16:0-DMA and 17:0-DMA FAME products (peaks in red) are also shown. Minor quantities of 18:0-DMA (0.1%) were also detected in *C. perkinsii.* Plasmalogen abundance (as DMAs) is calculated as a percentage of the total lipid content in the FAME GC-MS analysis. (**C**) Complementation analyses as in (A), with the PEDS1 homologs from *T. harzianum* (*Th*), *T. aggressivum* (*Tag*), *Thozetella* sp. (*Thz*), and *M. scopiformis* (*Ms*).

Consistent with our validation of PEDS1 homologs, plasmalogens have been reported for *T. cruzi, L. major* and *D. discoideum*, with C16 or C18 *sn*-1 vinyl ether-linked chains (21, 36, 37, 39, 44, 45, 53, 54) as is commonly observed in Metazoa (13, 14, 37, 47–50). Our FAME GC-MS analyses of *C. perkinsii* lipid extracts revealed the presence of plasmalogens with C16:0 vinyl ether-linked chains (detected as 16:0-DMA) (**Fig. 3B**), together with trace amounts (0.1%) of plasmalogens with C18:0 at the *sn*-1 position. This finding uncovers that the only Ichthyosporean with PEDS1 identified to date produces mainly plasmalogens with C16 at *sn*-1, a chain length that is common among eukaryotes. A very recent study (22) as well as our FAME GC-MS analysis (**Fig. 3B**) showed plasmalogens also in *C. owczarzaki*, but with predominantly C17:0 chains at the *sn*-1 position. Differences in chain lengths among species are likely related to substrate availability and species-specific requirements of FAR, GNPAT and/or AGPS (30, 55–57). Although complete lipidomics data are not available for *P. chileana* or *S. regulata*, the presence of AGPS and FARAT (or GNPAT and FAR as separate proteins) (**Fig. 2A**), together with a functional PEDS1, would suggest the capacity for plasmalogen biosynthesis in the two species.

Given the presence of PEDS1 homologs in a restricted number of Fungi (**Fig. 2B, Supplementary Fig. S2** and **Supplementary Table S6**), a group generally considered to lack plasmalogens, we also performed complementation analyses of the *M. xanthus* Δ*carF* strain with the fungal PEDS1 hits (all N-terminally FLAG-tagged) from (**Supplementary Fig. S4B** and **Supplementary Table S7**): *Trichoderma harzianum* (strain CBS 354.33), *Trichoderma aggressivum* (CBS 100526) and *Thozetella* sp. (PMI_491), where we identified PEDS1, AGPS and FARAT; and *M. scopiformis* (CBS 120377), where we could identify only PEDS1. MxVEPE production was observed with the *Thozetella* and *M. scopiformis* homologs, which could be detected by Western blot, though less than with *M. xanthus* CarF (**Fig. 3C**). The other two fungal homologs tested did not express well in *M. xanthus* (despite codon usage optimized for *M. xanthus*), likely explaining the negligible MxVEPE produced with *T. harzianum* and *T. aggressivum* homologs (**Fig. 3C**). Nonetheless, these results confirm for the first time PEDS1 activity for at least two fungal homologs, when expressed in *M. xanthus*. To obtain further experimental evidence, we examined the presence of plasmalogens by FAME GC-MS analysis in two fungal species with PEDS1, AGPS and FARAT homologs that were available from culture collections, *T. harzianum* (CBS 354.33) and *T. aggressivum* (CBS 100526), and whose PEDS1 was analyzed in *M. xanthus* (**Fig. 3C**) (see **Materials and Methods;** *Thozetella* sp.PMI_491, whose PEDS1 functioned best in *M. xanthus*, could not be obtained from culture collections). This analysis did not reveal any DMA species indicative of plasmalogens in either species. The presence of key biosynthetic enzymes yet no plasmalogens may be due to poor enzyme expression and/or activities, which could be addressed in future studies.

### Evolutionary Origin of CarF/PEDS1

Given the evident similarities between plasmalogen biosynthesis in eukaryotes and certain bacteria, specifically those in the phylum Myxococcota, we sought to determine the origin of this metabolic pathway in eukaryotes. For this, we searched the Genome Taxonomy Database (GTDB) (58), which provides a phylogenetically consistent and rank-normalized genome-based taxonomy for prokaryotes. We looked for PEDS1, AGPS, GNPAT, and FAR (or FARAT) using as queries the complete set of eukaryotic sequences identified in the previous section (see **Materials and Methods**). Although individual genes were detected across several bacterial phyla, only six phyla contained at least one genome with the complete gene set. Overall, we identified 73 bacterial genomes encoding the full pathway: 61 in the phylum Myxococcota, and the remaining 12 distributed across five additional phyla (**Supplementary Fig. S5** and **Supplementary Tables S1, S2, S4 and S5**). These findings indicate that the capability for synthesizing plasmalogens and their precursors aerobically is currently concentrated mostly within Myxococcota in bacteria.

To explore the evolutionary origins of the oxygen-dependent desaturase PEDS1, we performed phylogenetic analysis including all sequences retrieved from both prokaryotic and eukaryotic datasets. For eukaryotes, we incorporated all identified PEDS1 hits, encompassing both PEDS1-like and FAD4-like homologs (**Fig 2**). The phylogenetic reconstruction recovered the topology of the desaturase gene distribution revealing two distant monophyletic groups, one comprising the PEDS1-like homologs in Amorphea and Discoba and the other FAD4-like homologs in Diaphoretickes (**Fig. 4**).

**Figure 4.**
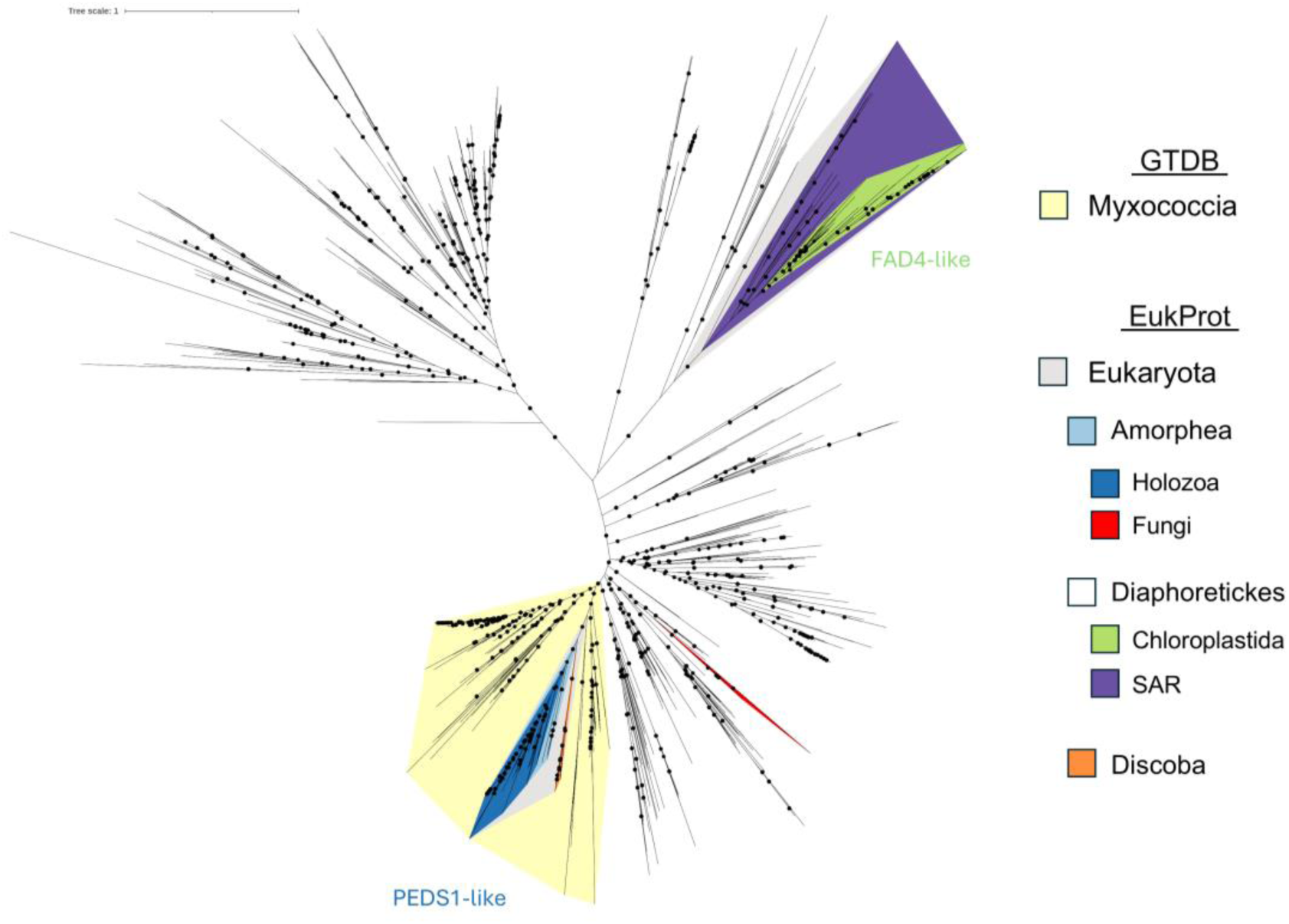
Phylogenetic tree for PEDS1 homologs in Bacteria and Eukaryotes. Maximum likelihood phylogenetic tree inferred using PEDS1 homologous protein sequences (137 amino acid positions) obtained from the GTDB and EukProt databases plus Fungi sequences retrieved by BLASTP NCBI. Colors correspond to relevant taxonomic classifications. For prokaryotes, the colors represent phylum Myxococcota. For eukaryotes, the represented clades are Amorphea (Holozoa; and Amoebozoa, as those sequences within Amorphea that do not correspond to Holozoa), Diaphoretickes (Chloroplastida, SAR) and Discoba. Holozoa within Amorphea and Chloroplastida or SAR within Diaphoretickes are highlighted. Black dots indicate ultrafast bootstrap (UFB) values greater than 75%.

Within the portion of the tree encompassing Amorphea and Discoba, the topology is consistent with known species relationships. First, all eukaryotes from these two clades form a monophyletic group (**Fig. 4**), which aligns with the presence/absence patterns described earlier (**Fig. 2A**). These are, in fact, the only major eukaryotic clades that harbour PEDS1. Within this monophyletic group, Holozoa and Amoebozoa (**Fig. 4**, **Supplementary Fig. S6** for details) each form distinct, closely related clusters, while a more basal lineage comprises Discoba. There are two exceptions: *N. gruberi*, which instead of clustering with Discoba appears on an independent branch between Amoebozoa and Holozoa, and *E. gracilis*, which is placed with the FAD4-like Diaphoretickes group (**Supplementary File 1**, for details). Notably, the entire Amorphea and Discoba monophyletic assemblage of eukaryotes is closely related to PEDS1/CarF from the Myxococcota phylum. Taken together, the observations suggest that an early eukaryote may have acquired PEDS1 through an early HGT event from Myxococcota. Moreover, fungal PEDS1 sequences retrieved from the NCBI were included in the phylogenetic analyses and they formed a monophyletic group separated from the other eukaryotes (**Fig. 4**). If the fungal PEDS1 had been derived from the same ancestral HGT event, the fungal PEDS1 proteins would have been expected to cluster at least near the PEDS1-like group, ideally closer to Holozoan sequences. However, the fungal group lies apart from the other eukaryotic PEDS1 and, instead, is more closely related to bacterial PEDS1 (**Supplementary Fig. S6**). This finding would suggest that the PEDS1-like sequences identified in Fungi arose from an independent HGT event.

Regarding FAD4-like proteins, the only monophyletic group we identified is Chloroplastida, with two noteworthy outliers: *Cyanophora paradoxa* (Glaucophyta) and, most strikingly, *E. gracilis*, which belongs to the supergroup Discoba and lacks PEDS1, as mentioned before. It should be noted that *E. gracilis* is photosynthetic, a trait acquired through a secondary endosymbiosis event involving a green alga (59). The well-documented nature of this endosymbiosis is consistent with *E. gracilis* FAD4 clustering alongside Ostreococcus and Micromonas sequences in the phylogenetic tree (**Supplementary Fig. S7**). The remaining organisms that harbor FAD4-like enzymes (Haptophyta, Cryptomonadales, Rhodophyta, Ochrophyta and Colpodellida) do not form clear monophyletic clusters, suggesting a more complex evolutionary history. Although beyond the scope of this study, it would be worthwhile examining whether these phylogenetically related proteins truly fulfil a FAD4 role.

### Evolutionary origin of the other enzymes of the plasmalogen biosynthesis pathway

We next examined the origin of the remaining enzymes in the pathway. Since the AGPS sequences did not provide sufficient phylogenetic signal, we focused on FAR and GNPAT that, as shown in this study, exist as a fused FARAT protein in most eukaryotic groups. A comprehensive phylogenetic analysis using all retrieved GNPAT sequences was first performed (see **Materials and Methods**) but since the dataset was extensive, redundancy was reduced by sequence clustering based on identity and coverage (see **Supplementary Table S8**). Unlike the distinct clades observed for PEDS1, our analysis of FARAT revealed a single monophyletic group encompassing all eukaryotes, including those in Discoba, Amorphea, and Diaphoretickes (the latter being represented solely by Ciliophora) (**Fig. 5, Supplementary Fig. S9** for details). In principle, this would imply a single evolutionary origin of FARAT in eukaryotes. However, since the relationships among subgroups deviate from those expected under strict vertical inheritance, a more intricate evolutionary history is suggested, as detailed below. The most basal eukaryotic clade is composed by Discoba, specifically by the Kinetoplastea and Euglenida, which display a topology consistent with known species relationships (60, 61). However, rather than Discoba, the Heterolobosea (*N. gruberi* and *N. damariscottae*) clustered close to Ciliophora, and both appeared more related to Amoebozoa (**Supplementary Fig. S8 and Supplementary File 2**). Since a similar pattern was observed for PEDS1 in *N. gruberi*, it could be that *N. gruberi* replaced its FARAT and PEDS1 through a secondary HGT event, possibly from an Amoebozoa. As for Ciliophora, lack of PEDS1 (for direct comparison with FARAT) precludes a more definitive assessment of this scenario; nonetheless, high bootstrap in the tree would support that FARAT originated in this group from an HGT event among eukaryotes (**Supplementary Fig. S8**), which would be consistent with Ciliophora being the only Diaphoretickes with FARAT (**Fig. 2A**). Given that the FARAT proteins from Ciliophora and Heterolobosea form sister groups, it is plausible that an HGT event occurred from an amoeba to one of these groups, which then acted as an intermediary for a subsequent HGT with its sister group.

**Figure 5.**
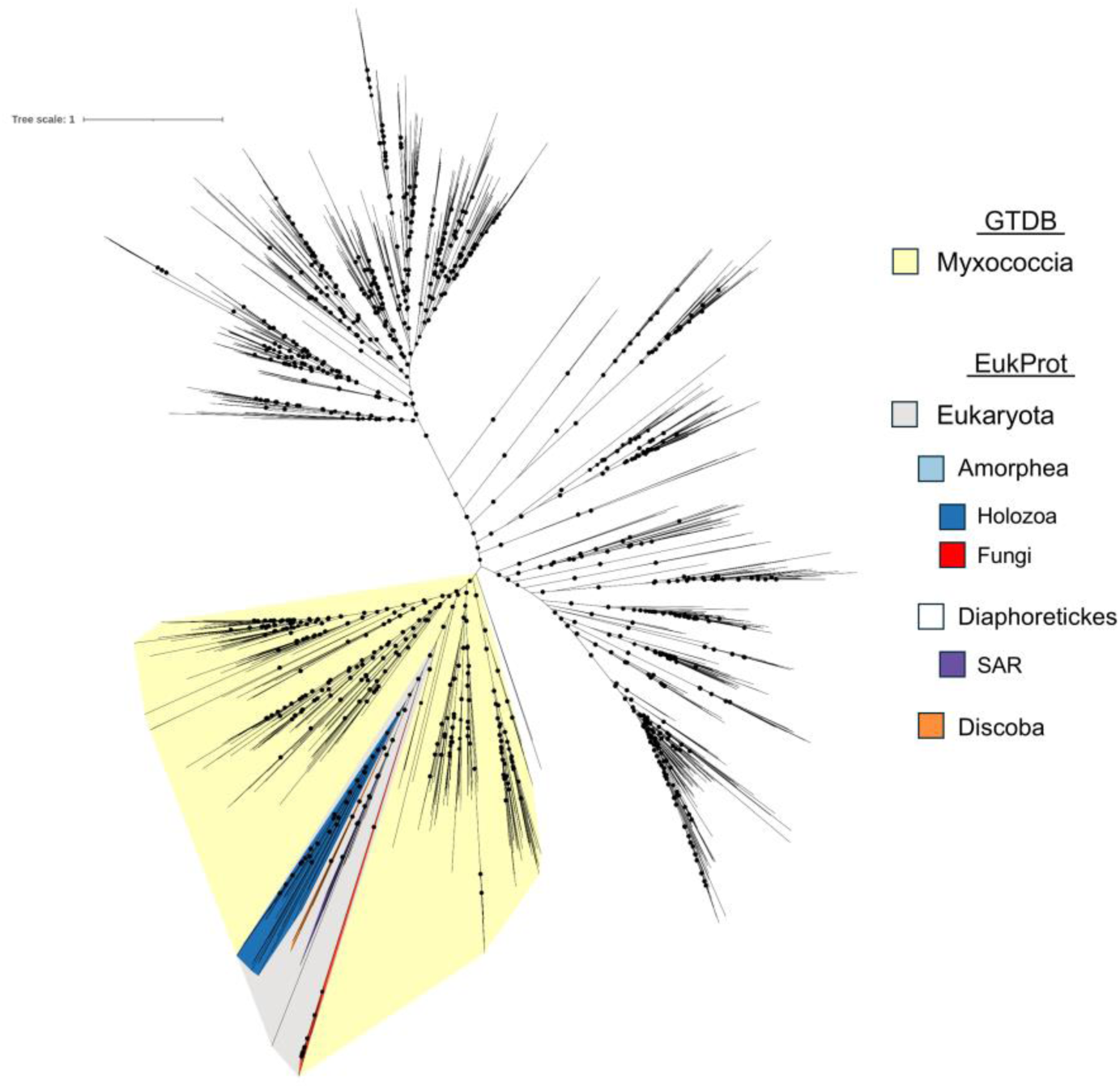
Phylogenetic tree illustrating FARAT homologs in Bacteria and Eukaryota. Maximum likelihood phylogenetic tree generated using GNPAT homologous protein sequences (960 amino acids positions) obtained from the GTDB and EukProt databases plus Fungi sequences retrieved by BLASTP NCBI. Colors correspond to relevant taxonomic classifications. For prokaryotes, the colors represent phylum Myxococcota. For eukaryotes, the represented clades are Amorphea, Discoba, and Diaphoretickes. Black dots indicate ultrafast bootstrap (UFB) values greater than 75%.

The eukaryotic FARAT proteins appeared closely related to sequences from Myxococcia, suggesting that FARAT was acquired via HGT from phylum Myxococcota, as was also proposed for PEDS1. Moreover, the phylogenetic tree supports a single HGT event also for FARAT, which likely occurred before the divergence of the Discoba and Amorphea lineages, given the monophyletic relationship and clade topology. As previously noted, while Amoebozoa, Discoba, and SAR possess FARAT, the separated GNPAT and FAR into distinct proteins appears to be exclusive to Holozoa (with some exceptions). Since these groups form a monophyletic clade, the most likely scenario would be the acquisition of the bifunctional gene from Myxococcota by an early eukaryote, followed by a fission event that gave rise to two independent genes now present in Holozoa, along with a few additional independent fission events in other lineages. Additionally, FAR appears to have undergone expansion in Holozoa, forming a gene family, an aspect that was not explored further in this study. FARAT sequences retrieved from the limited number of Fungi with PEDS1 did not follow the distribution expected for vertical inheritance. In fact, they formed a monophyletic group of Fungi located very basally within the eukaryotic branch (**Fig. 5, Supplementary Fig. S9** for details). This contrasts with what was observed for PEDS1, where fungal sequences formed a clear cluster separated from the other eukaryotes. Thus, it is more difficult to support the assertion of an independent HGT event to explain the presence of FARAT in a selected group of Fungi.

## Discussion

This study represents the most comprehensive taxonomic investigation to-date of genes essential for plasmalogen production across the tree of life. To minimize analytical biases and ensure robust phylogenetic inference, we employed comprehensive reference databases that encompass a wide range of biological diversity (33, 58). Overall, our study sheds light into a key event in the evolution of eukaryotic cell complexity and the role of HGT in tailoring metabolic capacities. Our results for PEDS1 homologs accords with the occurrence of two related but distinct types of lipid desaturases: PEDS1-like and FAD4-like. PEDS1-like enzymes were detected exclusively in Discoba and Amorphea, whereas FAD4-like ones were found restricted to Diaphoretickes, suggesting a pattern of vertical inheritance within each supergroup. Because PEDS1 is indispensable for plasmalogen production, this observation *per se* effectively divides eukaryotes into two major groups: those that may potentially synthesize plasmalogens and those that cannot. Consistently, the other defining genes of the aerobic plasmalogen biosynthesis pathway generally co-occur with PEDS1 (**Fig. 2**), underscoring the evolutionary importance for some groups of maintaining a complete plasmalogen biosynthetic pathway.

Of interest is the observation that PEDS1-like and FAD4-like lipid desaturases never co-occur in the same organism. While independent evolutionary trajectories might explain this mutually exclusive distribution, the case of *E. gracilis* within Discoba is particularly noteworthy. Thus, while most Discoba clades possess all the defining enzymes of the aerobic pathway, *E. gracilis* retains AGPS and FARAT but lacks PEDS1 and, instead, has FAD4. Studies in myxobacteria and various Metazoa firmly established that plasmalogen synthesis absolutely requires PEDS1 in the final indispensable step of vinyl ether bond formation (13, 14, 23, 24, 49, 50), clearly explaining why *E. gracilis* lacks plasmalogens but produces the plasmanyl GP precursors (37, 38, 43), as expected given the presence of AGPS and FARAT. FAD4 in *E. gracilis* likely originated from the green algal endosymbiont (59), consistent with our finding that FAD4 is prevalent among the Chloroplastida and with the knowledge that green algae, similar to land plants, produce the unique thylakoid membrane lipid whose biosynthesis requires FAD4 (41, 42). The observation of FAD4 and no PEDS1 in *E. gracilis* supports the hypothesis that the presence of both enzymes might be mutually exclusive, perhaps reflecting an incompatibility between their respective lipid products in cellular membranes. If this were the case, a similar replacement process as in *E. gracilis* may have occurred in Diaphoretickes. The striking absence of PEDS1 in photosynthetic organisms like *E. gracilis* could be related to the susceptibility of the plasmalogen vinyl ether bond to cleavage by reactive oxygen species like singlet oxygen (^1^O_2_), which in turn produces other reactive species that are toxic to the cells (14, 62–65). Thus, photosynthesis would exert a selective pressure against plasmalogens and PEDS1 because it generates ^1^O_2_ as a byproduct of chlorophyll photoexcitation and energy transfer to oxygen at much higher levels than do other cellular processes (66). Interestingly, a recent study reported the unexpected discovery of aerobic anoxygenic photosynthesis in some members of phylum Myxococcota (67), and our search in their genomes (BioProject ID PRJNA943119) failed to detect any CarF/PEDS1 homologs, consistent with absence of PEDS1 in photosynthetic organisms.

In the paradigmatic mammalian plasmalogen biosynthesis pathway, FAR and GNPAT exist as two separate enzymes acting in two parallel reactions that provide the substrates for AGPS to catalyze ether bond formation (**Fig. 1A**). Our present study highlights a possibly underappreciated fact, namely the widespread presence of FAR and GNPAT as a fused FARAT protein in Discoba, Amoebozoa, Filasterea (Obazoa), as well as in Ciliophora (SAR) where a FARAT was first reported (30); and, as with the standalone GNPAT and FAR, the bifunctional FARAT is also a peroxisomal enzyme (30, 45). Both FAR and GNPAT activities of FARAT occur within the peroxisome as does the monofunctional metazoan GNPAT (30, 45), but metazoan FAR, although associated with the peroxisomal membrane, appears to have its activity located on the cytosolic side (68). This suggests physical separation of the FAR and GNPAT activities in Metazoa, which might ensure tighter spatial control, while a bifunctional FARAT might enable a more efficiently coordinated regulation. Importantly, our findings suggest that eukaryotes originally acquired GNPAT and FAR as encoded by a single bifunctional gene that is retained as such in most groups, perhaps from a selective advantage in having both activities combined in one protein. The gene then underwent a fission event in the Holozoa branch (Obazoa; Opisthokonta) resulting in two separate genes that became subsequently fixed in Metazoa (a group that includes many model organisms) with loss of FARAT. Thus, the mammalian pathway architecture as regards FAR and GNPAT should be considered as the outcome of a more recent evolutionary event and as an exception rather than the norm.

A recent study proposed that PEDS1 was acquired by Filozoa, Excavata, Amoebozoa, and some small number of Fungi through four independent HGT events, each from an unknown donor (22). In contrast, our analysis, which benefits from broader taxon sampling and refined search strategies, supports a model in which PEDS1 was acquired by an early eukaryote through a single HGT event from phylum Myxococcota, followed by a more recent HGT event from the same bacterial phylum into a small group of Fungi, several of which are soil-dwelling, a trait shared with myxobacteria and known to promote HGT (69, 70). Moreover, our results with FAR and GNPAT strongly suggest that FARAT was also acquired by an early eukaryote through one HGT event from Myxococcota. The apparent absence of plasmalogens in the handful of fungi with PEDS1, AGPS and FARAT, suggesting poor enzyme activity and/or expression, can be reconciled with relatively recent HGT, with the genes acquired not yet fully assimilated and optimized for function. Although these fungal genes do not retain the high GC-content and operon-like arrangement seen in myxobacteria, the narrow phylogenetic spread of these genes among fungi, and their low intron density (generally, zero or just one) and genomic proximity would be consistent with recent HGT (**Supplementary Fig. S2** and **Supplementary Table S6)** (71). Given that similar scenarios of HGT from Myxococcota have been documented before (72–76), it is plausible that several genes or even the entire metabolic pathway was transferred at once from Myxococcota into an early eukaryote. Among current hypotheses for eukaryogenesis, the syntrophic model discusses the evolutionary relationship between myxobacteria and early eukaryotes (77). This model proposes a tripartite metabolic symbiosis involving a complex sulfate-reducing myxobacterial-like host, an endosymbiotic hydrogen-producing Asgard-like archaeon (future nucleus) and a sulfite-oxidizing facultative aerobe alphaproteobacterium (future mitochondrion) for the origin of the eukaryotic cell. A key aspect of the syntrophy model is that it invokes a bacterial (rather than an archaeal) host, which readily explains the shared nature of bacterial and eukaryotic phospholipids, whether in the plasma membrane or the endomembrane system (77). Recent studies have described the HGT from Myxococcota to eukaryotes of the pathway for the biosynthesis of steroids, which are fundamental lipids of the eukaryotic membrane (74–76). This aligns with our findings, since plasmalogen biosynthesis and steroid biosynthesis are both oxygen-dependent, with their regulation shown to be linked in mammals (78). These studies therefore support the proposal of an essential relationship between aerobic myxobacteria (previously classified within the deltaproteobacteria) and early eukaryotes. We did not find any evidence of the plasmalogen metabolic pathway in Archaea, suggesting that the HGT event would have occurred in the Last Eukaryotic Common Ancestor (LECA) or pre-LECA rather than directly from an archaeal lineage. This reinforces the hypothesis that the acquisition of the plasmalogen biosynthesis pathway was an innovation in early eukaryotic evolution, potentially conferring adaptive advantages that shaped membrane composition and function.

In our analysis, the patchy distribution of the pathway in eukaryotes is explained by several independent events of secondary loss of the plasmalogen synthesis genes, an evolutionary process relevant and frequent in eukaryotic evolution (79, 80). The most evident pattern is the complete loss of the pathway in anaerobic organisms, such as those in Metamonada and Archamoebae. Notably, Archamoebae, the sole anaerobic group within Amoebozoa, consistently lacks the genes for plasmalogen synthesis. Future studies may build upon our findings on gene loss and gain to further elucidate the functional role of plasmalogens in eukaryotes.

It is important to acknowledge that, like previous studies, our work still grapples with the unresolved root of the eukaryotic tree (34). The relationships among the supergroups Metamonada, Discoba, Amorphea and Diaphoretickes remain partially unresolved (34), which makes it challenging to draw further inferences. Several possibilities arise from the observation that the genes we studied are almost entirely restricted to Discoba and Amorphea. The simplest scenarios include either an HGT event to Amorphea and Discoba (if they are sister groups) or one involving the LECA followed by secondary loss in Diaphoretickes and Metamonada. We favor the latter scenario, drawing on recent phylogenomic analyses suggesting that excavate features represent the ancestral state of eukaryotes (35) (therefore Amorphea and Discoba would not be sister groups), the accumulating evidence of multiple HGT events from Myxococcota to LECA or pre-LECA, and the support from the syntrophy hypothesis (73, 74). While HGT events that introduce new functions are likely to be fixed by selection, these functions are often reversible and transient. Non-essential traits tend to be lost over evolutionary time when selective pressures shift (81). Horizontal gene transfers in eukaryotes are generally rare and short-lived, with many acquired genes providing only temporary benefits before being lost. However, the genes that are retained over long evolutionary periods, as observed in our study, are likely to be functionally significant or relevant (82).

## Materials and Methods

### Homology searches

We used the same workflow for all genes analyzed in this study (PEDS1/CarF, AGPS, GNPAT, FAR) to capture a broad range of biological diversity. The Genome Taxonomy Database (GTDB (58)) was used for prokaryotic genomes and EukProt v3 (33), which comprises 375 genomes, for eukaryotic genomes. Additionally, the genome of *Pigoraptor chileana*, a representative of Filasterea (PRJEB52884) was incorporated. We began our searches by performing Position-Specific Iterated BLAST (PSI-BLAST 2.12.0+) (83), using as queries a set of well-annotated eukaryotic proteins (**Supplementary File 3**) against the EukProt database, with parameters “-num_iterations 3 -inclusion_ethresh 1e-5” (**Supplementary File 3**). All proteins retrieved from this initial search were then used as queries to search the GTDB with the same strategy. To enhance phylogenetic information and exclude pseudogenes or aberrant sequences that might affect subsequent alignment and trimming, we scanned the resulting FASTA files from both searches using InterProScan v5.67-99.0 (84). We also filtered by protein length based on expected length and length distribution of PSI-BLASTP results. We retained only those sequences that contained representative domains with an e-value threshold of 1e-20. Specifically, we kept: PEDS1/CarF/FAD4 sequences with the PF10520 lipid desaturase domain and length between 150 and 350 aa; AGPS sequences with the PF01565 (FAD linked oxidase, N-terminal), PF02913 (FAD-binding oxidoreductase/transferase, type 4, C-terminal) and length between 500 and 700 aa, and PTHR46568 domains (alkyldihydroxyacetonephosphate synthase); GNPAT sequences with the glycerol-3-phosphate *O*-acyltransferase/dihydroxyacetone phosphate acyltransferase PTHR12563 domain and length smaller than 2000 aa; and FAR sequences with the fatty acyl-CoA reductase PTHR11011 domain and length bigger than 300 aa. Because the PTHR12563 domain and our length filtering includes both GNPAT and GPAT2, we conducted an additional phylogenetic analysis to distinguish between them and select only those that clustered to hits annotated as GNPAT. Briefly, we aligned the sequences using MAFFT v7.526 (linsi option) (85), trimmed the alignment with trimAl v1.5.rev0 (86) using default parameters and the “-automated1” option, and generated maximum likelihood phylogenies using IQ-TREE multicore version 2.0.7 (87). The search for PEDS1 homologs in Fungi was performed using BLASTP against the non-redundant protein sequences database at the National Center for Biotechnology Information, restricting the search to Fungi (NCBI taxid: 4751). The small number of fungal genomes with PEDS1 hits were then individually subjected to a BLASTP search for AGPS, FAR and GNPAT. Genome analyses revealed that the gene for PEDS1 was recurrently found, in a convergent orientation, next to that for AGPS, and generally close to the gene for FARAT (**Supplementary Fig. S2**). In some instances, this genomic arrangement helped to identify and manually curate some of the hits (**Supplementary Table S6**).

### Phylogenetic analyses

For each protein of interest, we constructed a phylogeny using the homologs identified in both EukProt and GTDB. For AGPS and FARAT, as the number of hits made phylogenetic reconstruction infeasible, we performed an initial clustering step based on sequence identity using MMseqs2 (88) with the parameters “-c 0.8 -e 1e-10 --min-seq-id 0.7”. The sequences were aligned using MAFFT v7.526 (85), specifically the linsi option for accurate alignment. The alignments were manually curated to remove poorly aligned sequences, realigned, and then trimmed using trimAl v1.5.rev0 (86) with default parameters and the options “-automated1” (PEDS1, AGPS and FAR) or “-gt 0.1” (FARAT). The resulting alignment was used to reconstruct the phylogeny with the maximum-likelihood method implemented in IQ-TREE 2.0.7 (87). ModelFinder selected the model according to the Bayesian Information Criterion LG+F+R10 for the three phylogenetic inferences (FAR excluded).

### Protein structure prediction and alignment

To properly classify GNPAT as GNPAT or FARAT in eukaryotes, protein structure predictions were performed for all eukaryotic GNPAT identified by PSI-BLAST using ColabFold version 1.5.5 (89) on a local server, employing the colabfold_batch functionality. Structural alignments to identify the FAR and GNPAT domains were carried out using the easy-search option of FoldSeek version 9.427df8a (90) and only hits with an e-value < 1e-5 were considered valid. Human FAR1 (Q8WVX9, UniProtKB/Swiss-Prot) and human GNPAT (O15228, UniProtKB/Swiss-Prot) protein structures were used as queries. Alignment visualizations were generated using ChimeraX (91).

### Cell growth conditions for *M. xanthus*, selected protists and fungi

*M. xanthus* was grown at 33 °C at 300 rpm in liquid CTT (1% Casitone, 10 mM Tris-hydrochloride, pH 7.6, 8 mM MgSO_4_, 1 mM phosphate buffer) medium supplemented, if required, with antibiotic (kanamycin or Km at 40 µg/mL) or inducer (0.5 mM vanillate) for conditional gene expression. CTT-1.5% agar supplemented with antibiotic, and inducer (if required) was used for growth at 33 °C on solid medium. Plates were grown in the dark or, when specified, under white light using fluorescent lamps (∼10 W/m^2^ intensity). To evaluate light-induced yellow-to-red colony color phenotype, 6 µL of exponentially growing cultures were spotted on CTT plates with inducer (0.5 mM vanillate) and grown for 48 h in the dark or the light. A tan or pale-yellow color in the light is typically exhibited by cells unable to synthesize carotenoids because of the sensitivity to light of DKxanthenes (92). To obtain samples for lipid analysis, a 10 mL CTT (with 40 µg/mL Km and 0.5 mM vanillate) culture of the corresponding *M. xanthus* strain was grown to an optical density at 550 nm (OD_550_) of 0.7-1, when cells were harvested by centrifugation at 15,000 *g* for 10 min at 4 °C and stored at -80 °C until further use.

*Capsaspora owczarzaki* was grown axenically at 23 °C in 25 cm^2^ (T25) culture flasks with vented caps, containing 10 mL of lipid-conditioned medium: 1% Bacto peptone, 1% yeast extract, 0.1% ribonucleic acid type VI from Torula yeast, 15 mg/L folic acid, 1 mg/L haemin, 10% lipid-depleted fetal bovine serum (lipid-depleted FBS), 2% phosphate buffer (18.1 g/L KH_2_PO_4_, 25 g/L Na_2_HPO_4_). Lipid-depleted FBS was obtained as described previously (93). Cells used to inoculate the cultures were harvested from a confluent culture in complete medium (that is, with FBS that was not lipid-depleted) by centrifugation at 2,000 *g* for 3 min at RT in a swinging bucket rotor. The pellet was washed with 10 mL sterile 1X PBS, and the process was repeated twice. After final resuspension in 1X PBS, cells were counted and an aliquot of 5x10^4^ cells was used to inoculate 10 mL of conditioned medium in a T25 flask. Cells were incubated for 4 days (80% confluence), scrapped, and an aliquot transferred to a new T25 flask with conditioned medium. This process was repeated twice, for a total of 3 transfers, before using the cells for lipid extraction, to minimize the carryover of lipids from the complete FBS in the original culture. *C. owczarzarkii* cultures obtained following the indicated procedure were incubated until confluence (3-4 days), harvested by centrifugation and cell pellets conserved at -80 °C until further processing. *C. perkinsii* cells grown axenically in YM medium (3 g/L yeast extract, 3 g/L malt extract, 5 g/L peptone, 10 g/L dextrose, 20 g/L NaCl) at 23 °C protected from light, as described elsewhere (94), were stored at -80 °C until further use. Fungal strains (*T. harzianum* CBS 354.33 and *T. aggressivum* f. Europaeum CBS 100526), were obtained from the Westerdijk Fungal Biodiversity Institute collection (https://wi.knaw.nl/Collection) and grown at 21 °C on oatmeal agar as recommended. Mycelia were harvested from colonies grown on the agar media using a sterile pipette tip, exudated using a filter paper, and stored at -80 °C until further lipid analysis.

### Complementation analysis in *M. xanthus*

*M. xanthus* functional complementation was tested as follows. The coding sequence of the gene, fused to an N-terminal FLAG tag, was cloned into plasmid pMR3679 for conditional gene expression from a vanillate-inducible promoter (**Supplementary Table S7**). Coding sequences for N-terminal FLAG-tagged CarF homologs from *C. owckzarzaki* (*Co*), *C. perkinsii* (*Cp*), *P. chileana (Pc*), *S. regulata* (*Sr*), *L. major* (*Lm*), *T. cruzi* (*Tc)*, *D. discoideum* (*Dd*), *T. harzianum* (*Th*), *T. aggressivum* (*Tag*), *Thozetella* sp. (*Thz*), and *M. scopiformis* (*Ms*) were optimized for codon usage in *M. xanthus* and synthesized by GenScript. Each construct was introduced by electroporation into *M. xanthus* Δ*carF* strain, where integration into the genome occurs by homologous recombination at a chromosomal site with no promoter activity (95). Functional complementation was tested by growing each of the resulting strains (**Supplementary Table S7**) in the presence of vanillate and analyzing: (a) the yellow-to-red colony change on CTT plates exposed to light; (b) the presence of plasmalogen and ether lipids by GC-MS (see below).

### Western (immunoblot) analysis

*M. xanthus* cells expressing FLAG-tagged CarF homologs were grown in 10 mL CTT supplemented with Km and 0.5 mM vanillate. At OD_550_ ∼0.6, 1 mL culture was pelleted and resuspended in 200 µL of 20% (w/v) trichloroacetic acid (TCA, Sigma-Aldrich) containing cOmplete EDTA-free protease inhibitor (Roche), incubated on ice for 5 min, followed by the addition of 400 µL of 5% (w/v) of the same solution. This solution was centrifuged at 15,000 *g* for 5 min at 4 °C and the supernatant was discarded. The precipitate was washed once with 1 mL of 0.1% TCA and finally resuspended in 70 µL of loading buffer (50 mM Tris-HCl, pH 6.8, 2% SDS, 100 mM β-mercaptoethanol, 10% glycerol and 0.1% bromophenol blue) and 40 µL of 1 M Tris-HCl, pH 8. From this mixture, 25 µL were boiled for 5 min and run in 12% SDS-PAGE. Transference was made using a BioRad Trans-Blot Turbo system and ImmobilonP^SQ^ membranes (Millipore). The membrane was incubated with anti-FLAG® M2 antibodies (cat. no. F3165; Sigma-Aldrich) and, as loading control, the same blots were probed with anti-RNAP β (clone 8RB13, cat. no. 11544932; Thermo Fisher Scientific) monoclonal antibodies using ECL^TM^ Prime Western Blotting system (GE Health Sciences). Results were checked at least thrice for reproducibility.

### FAME GC-MS analysis

Total lipid was extracted and analyzed, as described previously (14). Briefly, cell pellets were transferred to Pyrex glass tubes, resuspended in 8 mL of chloroform:methanol (2:1 v/v) with vortexing, and then incubated for 1 h in ice. After mixing with 2 mL of 0.12 M KCl, 10 s vortexing, and centrifugation (400 *g*, 4 °C), the upper aqueous/protein phase was aspirated. The lower phase was filtered using a Whatman No. 1 filter paper, poured into a fresh pre-weighed Pyrex glass tube, dried under N_2_ stream at 35 °C, and weighed to determine the total lipid mass. Then, the dried lipids were resuspended in a mixture of chloroform:methanol (2:1 v/v) to a final concentration of 10 mg/mL, with 2,6-di-tert-butyl-4-methylphenol (BHT; Scharlab), at a final concentration of 0.01%, to minimize oxidation. To obtain the FAMEs, 1.5 mg of total lipids was dried with N_2_ at 35 °C and subjected to acid hydrolysis overnight at 55 °C with 1 mL toluene and 2 mL of 1% H_2_SO_4_ in methanol. The reaction was quenched with 2 mL 0.2 M KHCO_3_, mixed gently with 5 mL hexane:diethyl ether (1:1 v/v) containing 0.01% BHT, and then centrifuged for 2 min at 500 *g*. The upper hexane:ether phase was transferred to a separate tube, the lower phase was mixed again with 5 mL hexane:diethylether (1:1 v/v) and the upper phase transferred, combined with that of the earlier extraction, and dried with N_2_ at 35 °C. Dried samples were resuspended in 325 µL hexane and 75 µL of N-methyl-N-trimethylsilyltrifluoroacetamide (MSTFA) (Sigma-Aldrich), then incubated at 37 °C for 1 h. An aliquot of 100 µL of the final mixture were transferred to 1.5 mL SureStop amber vials (Thermo Fisher Scientific) for GC-MS analysis to analyze ether lipid and plasmalogen-derived OAGs and DMAs, respectively. An Agilent 7890B system with an HP-5ms (30 m x 0.25 mm, 0.25 µm) column and a 5977B electron impact mass selective detector with helium carrier gas at a flow rate of 1 mL/min were used for GC-MS analysis. The sample (2 µL) was injected in splitless mode, and the column temperature was ramped at 6 °C /min from 60 °C to 270 °C, where it was held for 10 min. Other settings were 250 °C, 280 °C, 230 °C and 150 °C, respectively, for the inlet, MSD transfer line, ion source and quadrupole temperatures; a fixed electron energy of 70 eV; and scan mode of the mass selective detector for a 40 to 600 m/z range. Data analysis of three independent replicates was carried out using Agilent MSD ChemStation (v. F.01.00.1903) software. For complementation assays, the relative abundance of i15:0-DMA is represented as the mean and standard deviation from three biological replicates, normalized to wild-type levels set at 100%. In the FAME GC-MS analysis of *C. owczarzaki* and *C. perkinsii*, plasmalogen abundance was calculated as a percentage of the total lipid content.

## Data, Materials, and Software Availability

All data in the study are included in the article and/or supporting information.

## Acknowledgments

This work was supported by grants PID2021-123336NB-C21 and PID2024-158644NB-C21 (to M.E.-A.); PID2021-123336NB-C22 and PID2024-158644NB-C22 (to S.P.); PID2023-153273NB-I00 (to I.R-.T.) from the Ministerio de Ciencia e Innovación (MCIN) and Ministerio de Ciencia, Innovación, y Universidades (MICIU)/Agencia Estatal de Investigación (AEI)-Spain (MCIN/AEI/10.13039/501100011033) and “ERDF A way of making Europe”; and grant 21939/PI/22 (to M.E.-A.) from Fundación Séneca-Murcia (Spain). I.R.-N was supported by a Ph.D. fellowship (PRE-2019-090944) from the MCIN/AEI, and J.M.T.-B. was supported by a scholarship from Comisión Académica de Posgrado, Universidad de la República (cap.posgrados.udelar.edu.uy).

## Notes

**Competing Interest Statement:** The authors have no competing interests.

### Competing Interest Statement

The authors have declared no competing interest.

