## Supplementary Figures for "Origin of Eukaryotic Plasmalogen Biosynthesis by Horizontal Gene Transfer from Myxobacteria"

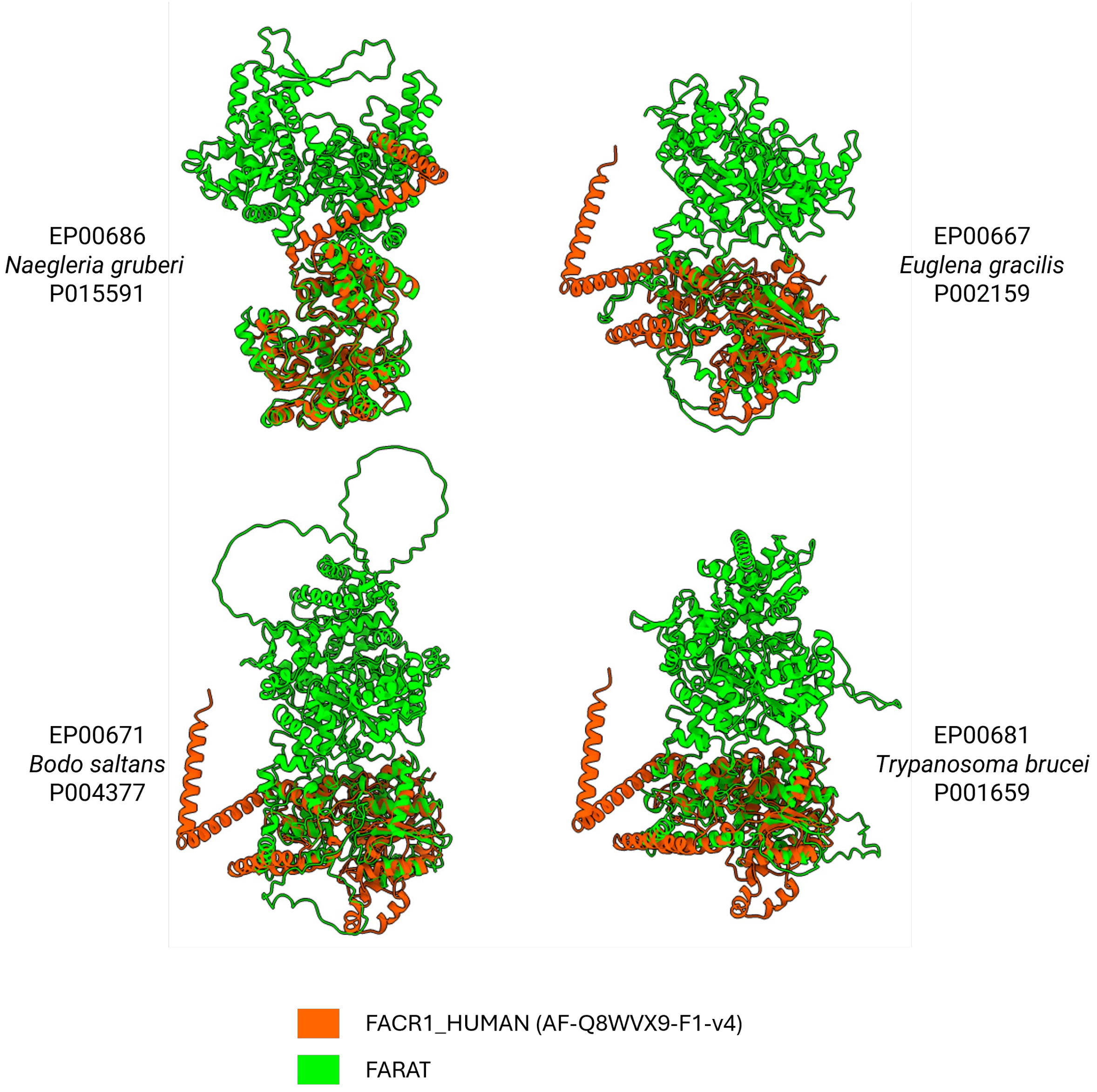


**Supplementary Figure S1. Structural alignment of Discoba FARAT proteins.** Structural alignment of putative FARAT proteins (green) from four Discoba species: *Naegleria gruberi* (EP00686), *Euglena gracilis* (EP00667), *Bodo saltans* (EP00671) and *Trypanosoma brucei* (EP00681) with human FAR protein (orange) (UniProt ID: FACR1_HUMAN, AF-Q8WXV9-F1-v4). The TM-scores indicate the degree of structural similarity (higher values suggest greater structural alignment): *N. gruberi* (0.87), *E. gracilis* (0.49), *B. saltans* (0.46) and *T. brucei* (0.51). Additional details on the structural alignments are provided in Supplementary Table S3.

FIGURE S1

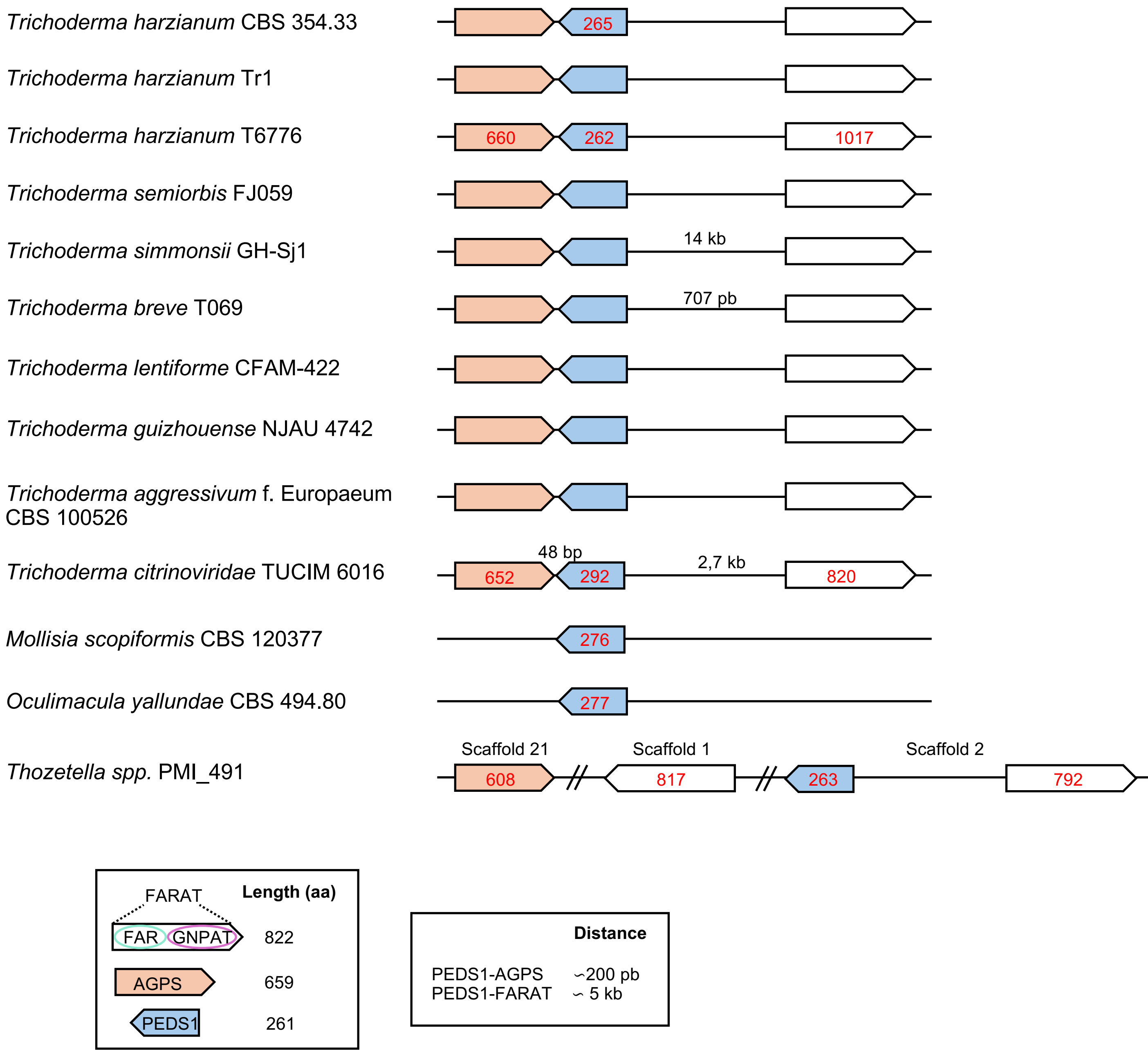


**Supplementary Figure S2. Genomic proximity of genes for the plasmalogen biosynthesis enzymes among Fungi with PEDS1.** Fungal species with homologs to PEDS1, AGPS, GNPAT and FAR (the latter two fused as FARAT) and schematic showing genomic proximity of the corresponding genes. Numbers in red within the genes (represented as arrows), which indicate the size (in amino acids) of the corresponding gene products, are included if the size differs from that found in most cases (bottom left). Same applies to the intergenic distances denoted in black (nucleotides) (bottom right: distances found in most cases).

FIGURE S2

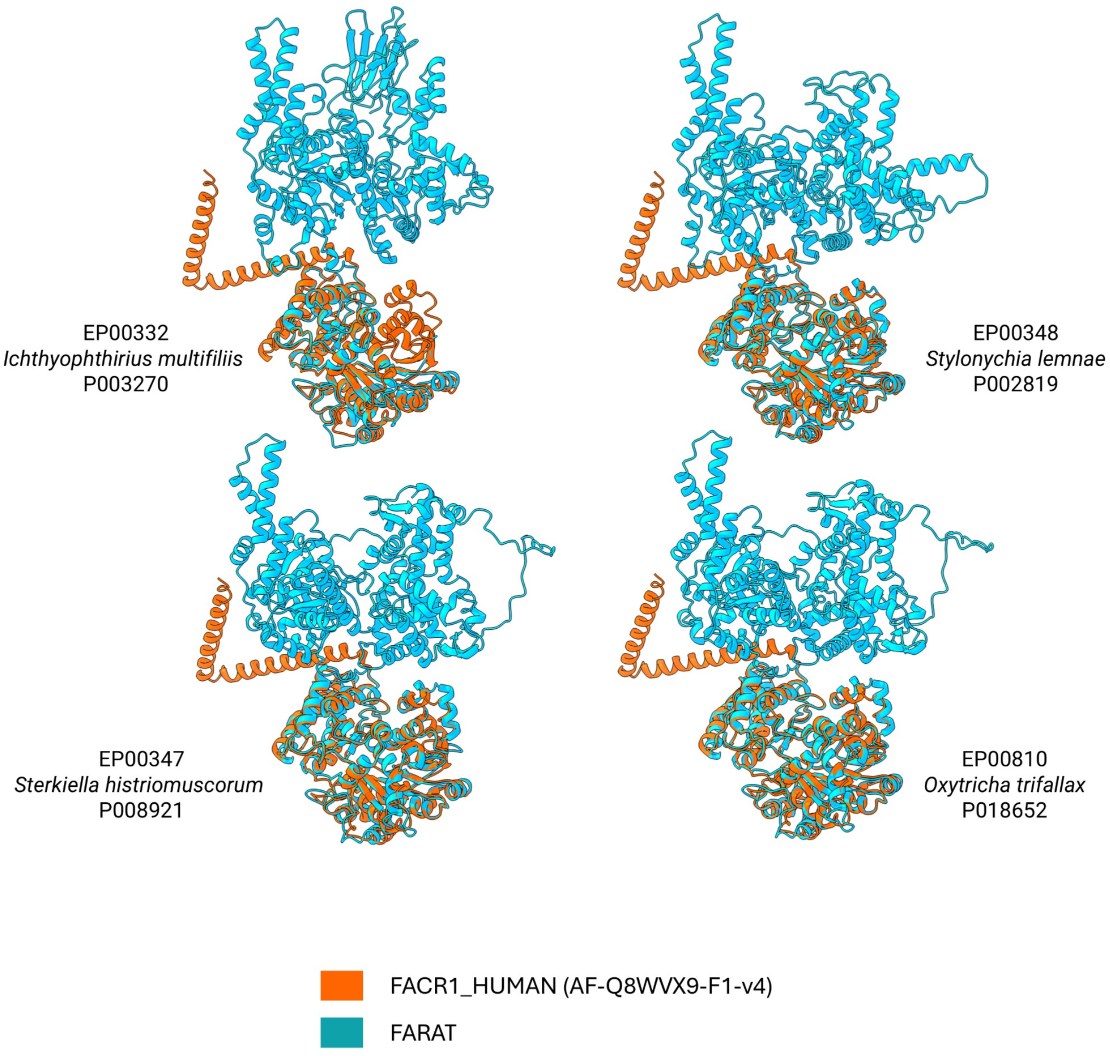


**Supplementary Figure S3. Structural alignment of Ciliophora FARAT proteins.** Structural alignment of putative FARAT proteins (blue) from four Ciliophora species: *Ichthyophthirius multifiliis* (EP00332), *Stylonychia lemnae* (EP00348), *Sterkiella histriomuscorum* (EP00347) and *Oxytricha trifallax* (EP00810) with the human FAR protein (orange) (UniProt ID: FACR1_HUMAN, AF-Q8WXV9-F1-v4). The TM-scores, indicate the degree of structural similarity (with higher values indicating greater structural alignment), are as follows*: I. multifiliis* (0.92), *S. lemnae* (0.92), *S. histriomuscorum* (0.89) and *O. trifallax* (0.89). Additional details on the structural alignments are provided in Supplementary Table S3.

FIGURE S3

A

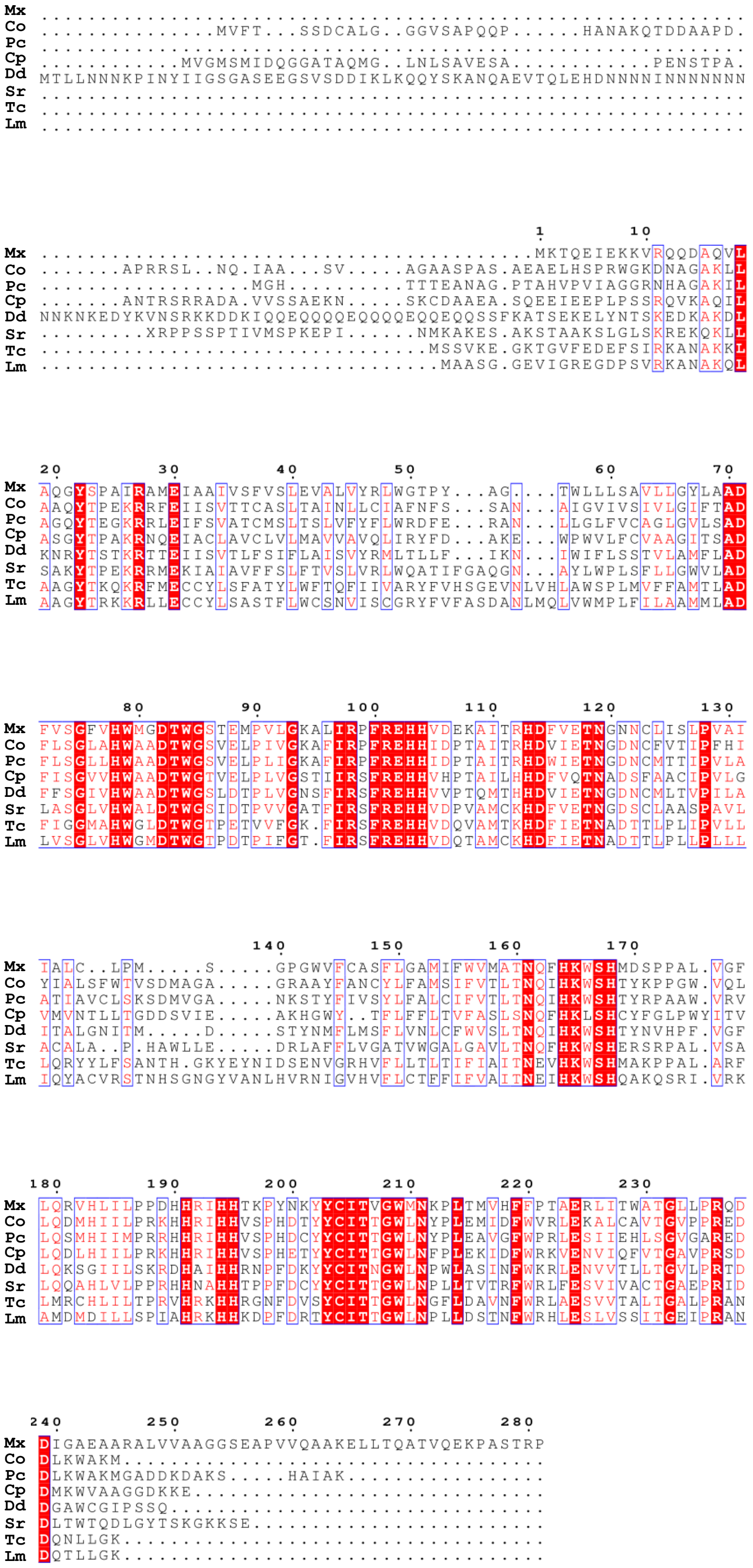
Mx ............................................................
Co ...............MVFT...SSDCALG..GGVSAPQQP......HANAKQTDDAAPD.
Pc ............................................................
Cp ............MVGMSMIDQGGATAQMG..LNLSAVESA............PENSTPA.
Dd MTLLNNNKPINYIIGSGASEEGSVSDDIKLKQQYSKANQAEVTQLEHDNNNNINNNNNNN
Sr ............................................................
Tc ............................................................
Lm ............................................................
1 10
Mx ..........................................MKTQEIEKKVRQQDAQVL
Co .......APRRSL..NQ.IAA...SV.....AGAASPAS.AEAELHSPRWGKDNAGAKLL
Pc ..................MGH..........TTTEANAG.PTAHVPVIAGGRNHAGAKIL
Cp .......ANTRSRRADA.VVSSAEKN.....SKCDAAEA.SQEEIEEPLPSSRQVKAQIL
Dd NNKNKEDYKVNSRKKDDKIQQEQQQQEQQQQEQQEQQSSFKATSEKELYNTSKEDKAKDL
Sr .........XRPPSSPTIVMSPKEPI......NMKAKES.AKSTAAKSLGLSKREKQKLL
Tc .................................MSSVKE.GKTGVFEDEFSIRKANAKKL
Lm ..................................MAASG.GEVIGREGDPSVRKANAKQL
20 30 40 50 60 70
Mx AQGYSPAIRAMEIAAIVSFVSLEVALVYRLWGTPY...AG....TWLLLSAVLLGYLAAD
Co AAQYTPEKRRFEIISVTTCASLTAINLLCIAFNFS...SAN...AIGVIVSIVLGIFTAD
Pc AGQYTEGKRRLEIFSVATCMSLTSLVFYFLWRDFE...RAN...LLGLFVCAGLGVLSAD
Cp ASGYTPAKRNQEIACLAVCLVLMAVVAVQLIRYFD...AKE...WPWVLFCVAAGITSAD
Dd KNRYTSTKRTTEIISVTLFSIFLAISVYRMLTLLF...IKN...IWIFLSSTVLAMFLAD
Sr SAKYTPEKRRMEKIAIAVFFSLFTVSLVRLWQATIFGAQGN...AYLWPLSFLLGWVLAD
Tc AAGYTKQKRFMECCYLSFATYLWFTQFIIVARYFVHSGEVNLVHLAWSPLMVFFAMTLAD
Lm AAGYTRKKRLLECCYLSASTFLWCSNVISCGRYFVFASDANLMQLVWMPLFILAAMMLAD
80 90 100 110 120 130
Mx FVSGFVHWMGDTWGSTEMPVLGKALIRPFREHHVDEKAITRHDFVETNGNNCLISLPVAI
Co FLSGLAHWAADTWGSVELPIVGKAFIRPFREHHIDPTAITRHDVIETNGDNCFVTIPFHI
Pc FLSGLLHWAADTWGSVELPLIGKAFIRPFREHHIDPTAITRHDWIETNGDNCMTTIPVLA
Cp FISGVVHWAADTWGTVELPLVGSTIIRSFREHHVHPTAILHHDFVQTNADSFAACIPVLG
Dd FFSGIVHWAADTWGSLDTPLVGNSFIRSFREHHVVPTQMTHHDVIETNGDNCMLTVPILA
Sr LASGLVHWALDTWGSIDTPVVGATFIRSFREHHVDPVAMCKHDFVETNGDSCLAASPAVL
Tc FIGGMAHWGLDTWGTPETVVFGK.FIRSFREHHVDQVAMTKHDFIETNADTTLPLIPVLL
Lm LVSGLVHWGMDTWGTPDTPIFGT.FIRSFREHHVDQTAMCKHDFIETNADTTLPLLPLLL
140 150 160 170
Mx IALC..LPM.....S.....GPGWVFCASFLGAMIFWVMATNQFHKWSHMDSPPAL.VGF
Co YIALSFWTVSDMAGA.....GRAAYFANCYLFAMSIFVTLTNQIHKWSHTYKPPGW.VQL
Pc ATIAVCLSKSDMVGA.....NKSTYFIVSYLFALCIFVTLTNQIHKWSHTYRPAAW.VRV
Cp VMVNTLLTGDDSVIE.....AKHGWY..TFLFFLTVFASLSNQFHKLSHCYFGLPWYITV
Dd ITALGNITM.....D.....STYNMFLMSFLVNLCFWVSLTNQIHKWSHTYNVHPF.VGF
Sr ACALA..P.HAWLLE.....DRLAFFLVGATVWGALGAVLTNQFHKWSHERSRPAL.VSA
Tc LQRYYLFSANTH.GKYEYNIDSENVGRHVFLLTLTIFIAITNEVHKWSHMAKPPAL.ARF
Lm IQYACVRSTNHSGNGYVANLHVRNIGVHVFLCTFFIFVAITNEIHKWSHQAKQSRI.VRK
180 190 200 210 220 230
Mx LQRVHLILPPDHHRIHHTKPYNKYYCITVGWMNKPLTMVHFFPTAERLITWATGLLPRQD
Co LQDMHIILPRKHHRIHHVSPHDTYYCITTGWLNYPLEMIDFWVRLEKALCAVTGVPPRED
Pc LQSMHIIMPRRHHRIHHVSPHDCYYCITTGWLNYPLEAVGFWPRLESIIEHLSGVGARED
Cp LQDLHIILPRKHHRIHHVSPHETYYCITTGWLNFPLEKIDFWRKVENVIQFVTGAVPRSD
Dd LQKSGIILSKRDHAIHHRNPFDKYYCITNGWLNPWLASINFWKRLENVVTLLTGVLPRTD
Sr LQQAHLVLPPRHHNAHHTPPFDCYYCITTGWLNPLLTVTRFWRLFESVIVACTGAEPRID
Tc LMRCHLILTPRVHRKHHRGNFDVSYCITTGWLNGFLDAVNFWRLAESVVTALTGALPRAN
Lm AMDMDILLSPIAHRKHHKDPFDRTYCITTGWLNPLLDSTNFWRHLESLVSSITGEIPRAN
240 250 260 270 280
Mx DIGAEAARALVVAAGGSEAPVVQAAKELLTQATVQEKPASTRP
Co DLKWAKM....................................
Pc DLKWAKMGADDKDAKS.....HAIAK.................
Cp DMKWVAAGGDKKE..............................
Dd DGAWCGIPSSQ................................
Sr DLTWTQDLGYTSKGKKSE.........................
Tc DQNLLGK....................................
Lm DQTLLGK....................................

FIGURE S4A

**B**

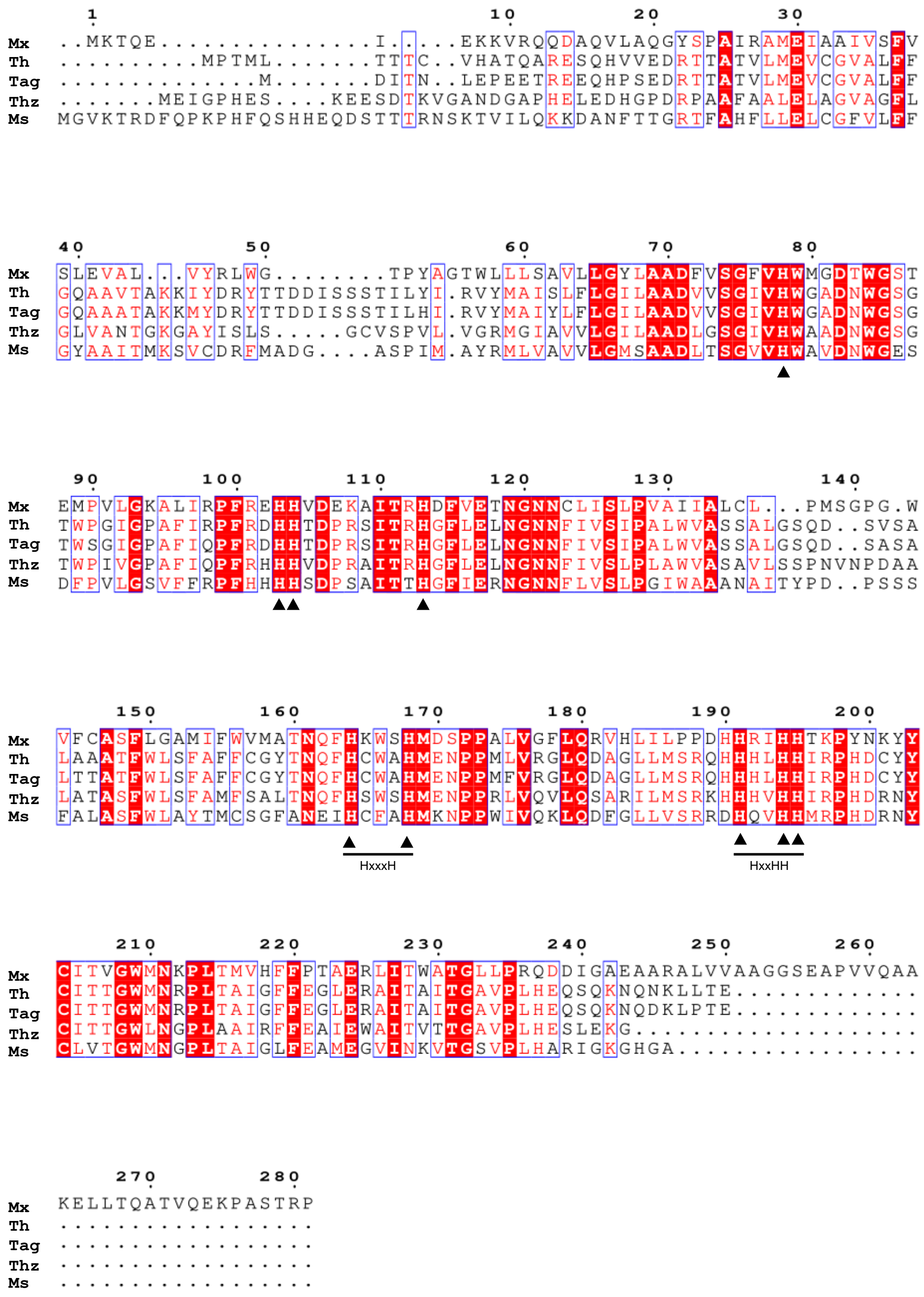


**Supplementary Figure S4. Sequence alignment of *M. xanthus* CarF and PEDS1/CarF homologs tested in complementation analyses.** (**A**) *M. xanthus* CarF and tested eukaryotic homologs from Filasterea (*Capsaspora owczarzaki* (Co) and *Pigoraptor chileana* (Pc)), Ichtyosporea (*Chromosphaera perkinsii* (Cp)), Amoebozoa (*Dictyostelium discoideum* (Dd) and *Stygamoeba regulata* (Sr)) and Kinetoplastea (*Trypanosoma cruzi* (Tc) and *Leishmania major* (Lm)). (**B**) *M. xanthus* CarF and tested homologs from Fungi: *Trichoderma harzianum* (Th), *Trichoderma aggressivum* f. Europaeum (Tag), *Thozetella* sp. (Thz), *Mollisia scopiformis* (Ms). In both sequence alignments: identical residues are shaded red; partially conserved residues are colored in red; histidine residues conserved in all sequences are indicated at the bottom by dark triangles; HxxxH and HxxHH motifs are also indicated.

FIGURE S4B

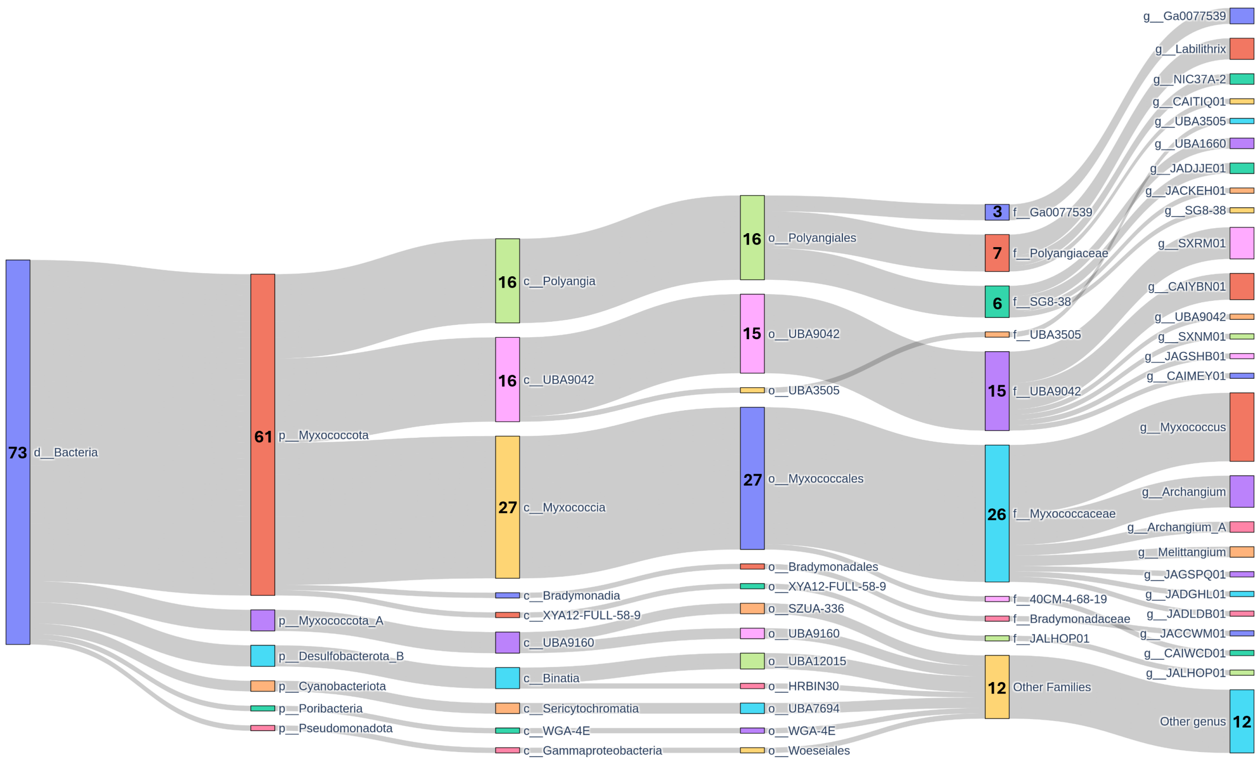


**Supplementary Figure S5. Sankey diagram of bacterial genomes harboring genes encoding possible PEDS1, AGPS, FAR and GNPAT activities.** This Sankeyplot displays all bacterial genomes identified as carrying at least one copy of each gene for the plasmalogen biosynthesis pathway. The diagram follows the GTDB-based taxonomic classification from domain (left) to genus (right). The numbers within each node represent the number of genomes falling into that particular category.

FIGURE S5

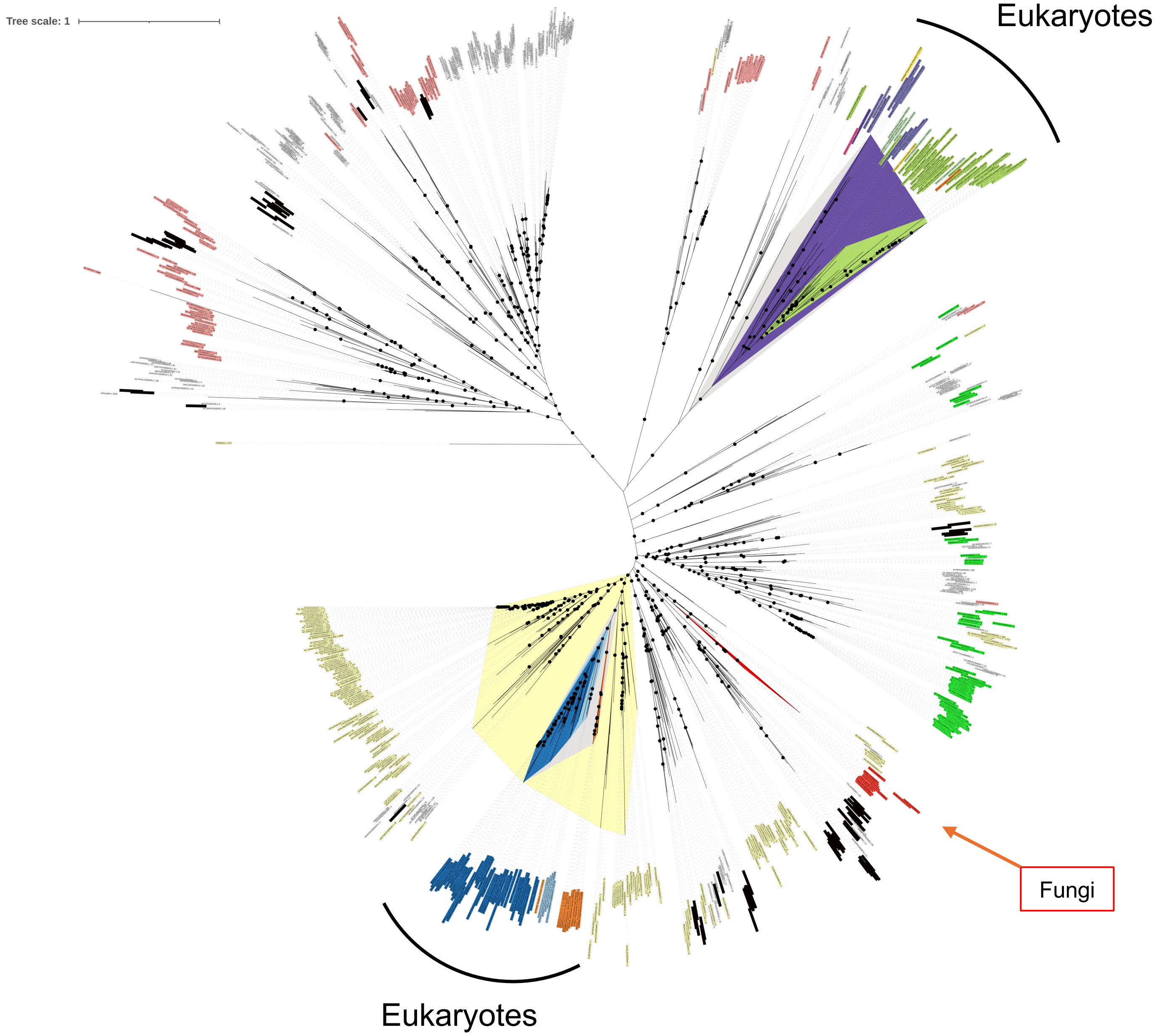


**Supplementary Figure S6. Phylogenetic tree for PEDS1 homologs in Bacteria and Eukaryotes.** Maximum likelihood phylogeny was generated using PEDS1 homologous protein sequences in the GTDB and EukProt databases plus Fungi sequences retrieved by BLASTP NCBI. Leafs are color-coded based on their taxonomic classifications as in Figures 2, 4 and 5. For most abundant prokaryotes phylums, the colors represent Myxococcota / Myxococcota_A (light yellow); Pseudomonadota (light peach/coral); Planctomycetota (very dark brown / near-black); Spirochaetota (bright green); Verrucomicrobiota (bright blue/violet). Black dots indicate ultrafast bootstrap values greater than 75%.

FIGURE S6

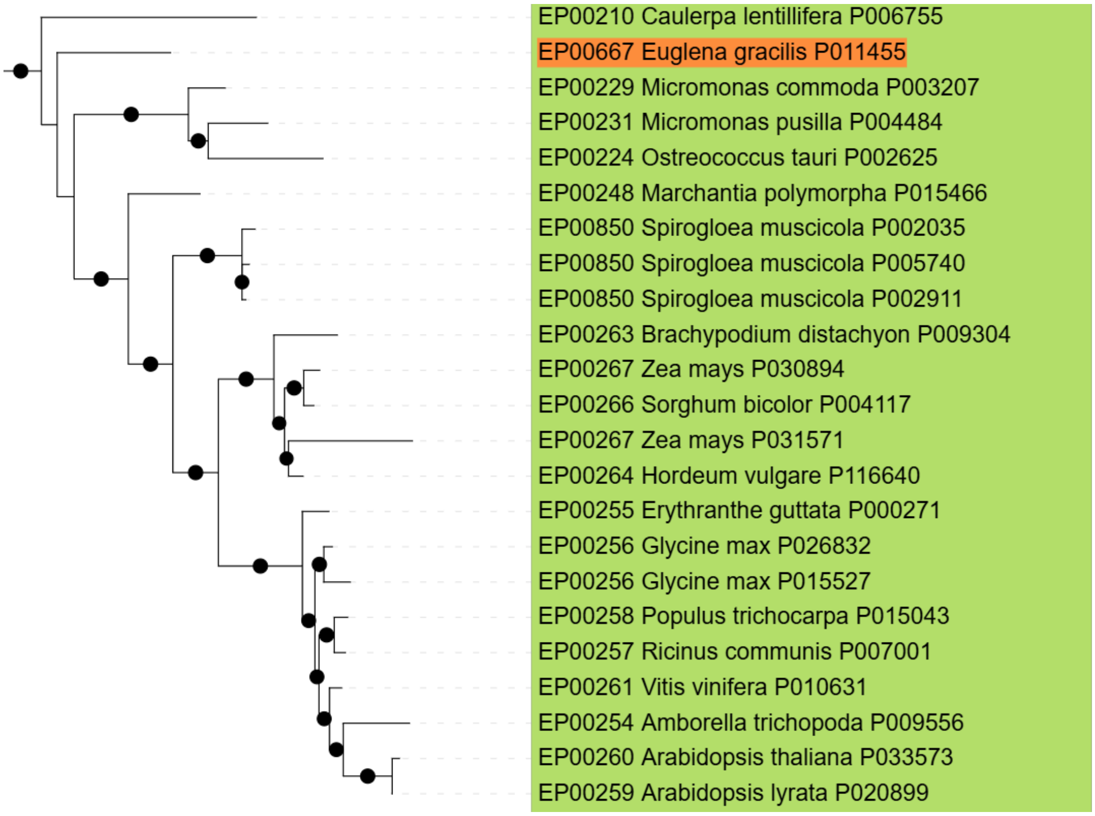


**Supplementary Figure S7. Phylogeny of *E. gracilis* FAD4-like protein.** Focused view of the portion of the main phylogenetic tree (Figure 4) that comprises FAD4-like lipid desaturase sequences. Branch support is indicated by black dots for ultrafast bootstrap values above 75%. Sequences are color-coded based on their taxonomic classifications as in Figures 2, 4 and 5.

FIGURE S7

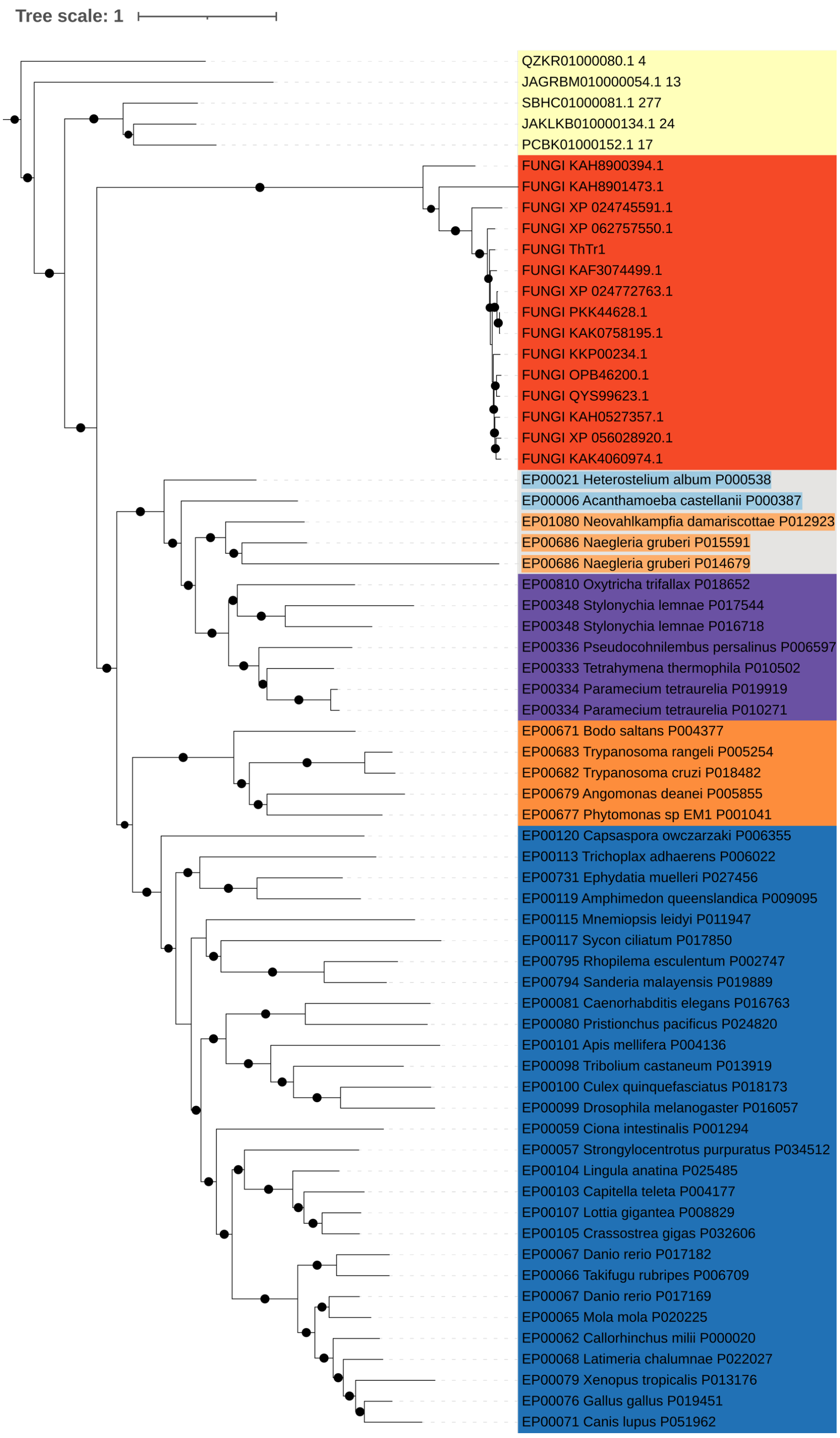


**Supplementary Figure S8. Phylogeny of FARAT proteins.** Focused view of the portion of the main phylogenetic tree (Figure 5) that comprises eukaryotic sequences. Branch support is indicated by black dots for ultrafast bootstrap values above 75%. Sequences are color-coded based on their taxonomic classifications as in Figures 2, 4 and 5.

FIGURE S8

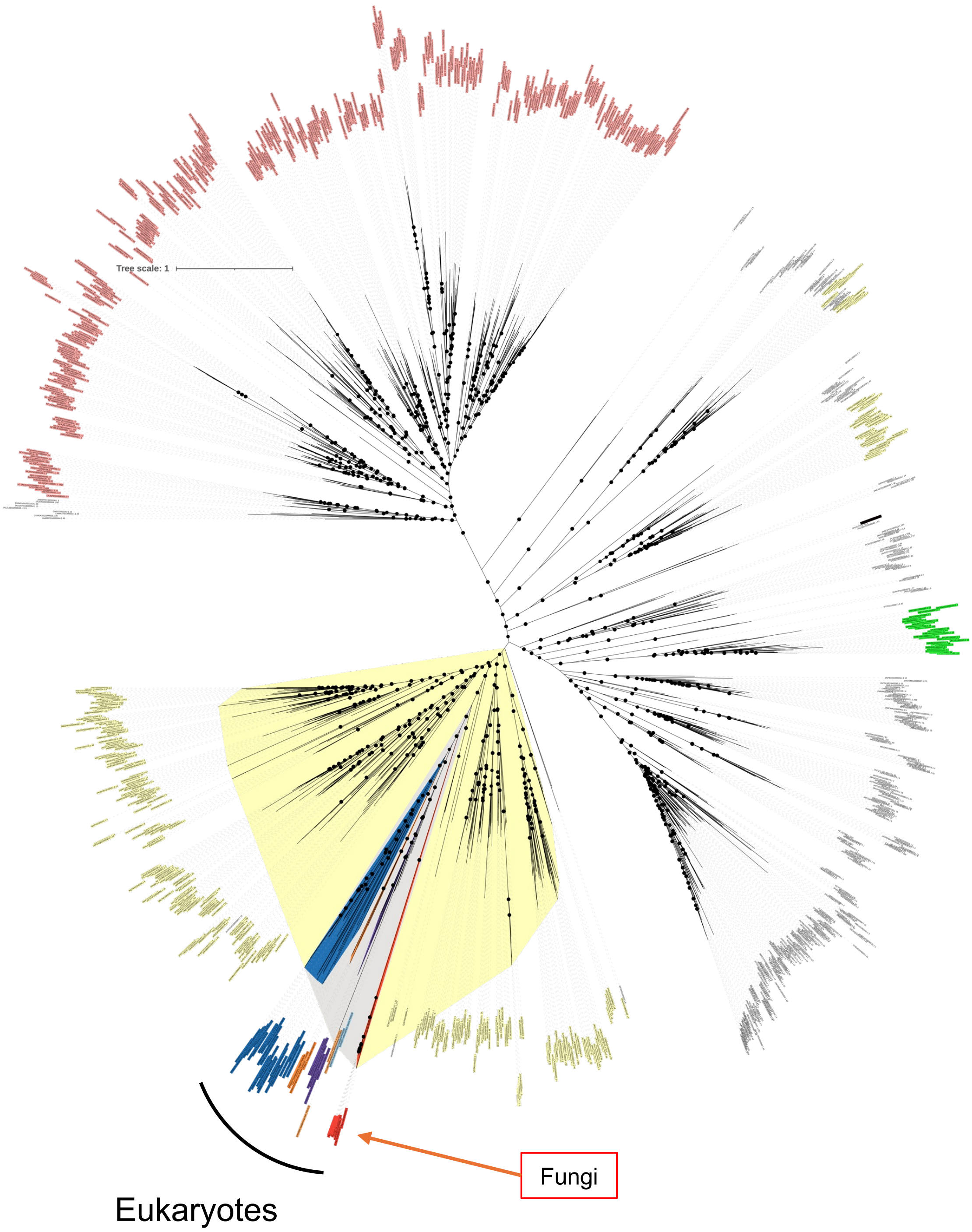


**Supplementary Figure S9. Phylogenetic tree for FARAT homologs in Bacteria and Eukaryota including Fungi sequences.** Maximum likelihood phylogeny was generated using GNPAT homologous protein sequences in the GTDB and EukProt databases plus Fungi sequences retrieved by BLASTP search at NCBI. Sequences are color-coded based on their taxonomic classifications as in Figures 2, 4 and 5. For most abundant prokaryotes phylums, the colors represent Myxococcota / Myxococcota_A (light yellow); Pseudomonadota (light peach/coral); Planctomycetota (very dark brown / near-black); Spirochaetota (bright green); Verrucomicrobiota (bright blue/violet). Black dots indicate ultrafast bootstrap values greater than 75%.

FIGURE S9
